# Computationally hybridized pathogenic mammarenavirus receptor binding domains reveal cryptic variants that elicit cross-neutralizing immune responses

**DOI:** 10.64898/2026.06.07.730625

**Authors:** Lily J. Taylor, Gavriella Siman-Tov, Sol Ferrero, Takeshi Saito, Gustavo Helguera, Junki Maruyama, Jose A. Rodriguez

## Abstract

The pathogenic mammarenaviruses, Machupo (MACV) and Junin (JUNV), are under evolutionary pressure to leverage human transferrin receptor 1 (hTfR1) for cellular entry while evading host immune responses during zoonosis. We now structurally and functionally investigate cryptic, computationally hybridized JUNV-MACV GP1 sequence variants. We then evaluate the ability of those variants to facilitate internalization of pseudotyped virus-like particles (PVs) into human cells and their recognition by neutralizing antibodies, including plasma from JUNV convalescent patients. We further compare the cryoEM structures of hTfR1-bound MACV GP1 to those of two functional hTfR1-bound MACV-JUNV hybrid GP1 variants that enable robust PV internalization and are also recognized by cross-neutralizing antibodies. Immunization with these variants demonstrates they and other hybrid GP1 sequences can elicit cross-reactive, neutralizing antibodies, supporting a model in which sequence adaptation within GP1 balances receptor recognition with immune evasion. This may inform the rational design of broadly neutralizing GP1-targeted antiviral therapies.

## Introduction

Mammarenaviruses are enveloped, bisegmented RNA-containing members of the *Arenaviridae* family, and are grouped geographically into Old World and New World species.^1^ New World mammarenaviruses (NWM) are divided into clades: A, B, A/B, and C; Clade B contains human pathogens: Sabia (SABV), Guanarito (GTOV), Chapare (CHPV), Machupo (MACV), and Junin (JUNV).^2^ This Clade also contains closely related non-pathogenic variants Tacaribe (TCRV), Amapari (AMAV), and Cupixi (CPXV).^3^ Each virus exists as a quasi-species, with a small number of well-annotated sequences, but a larger number of potential cryptic variants are expected to remain uncharacterized within their zoonotic hosts or to arise upon zoonosis.

New World mammarenaviruses (NWMs) contain a negative-stranded ambisense genome composed of a large (L) and small (S) segment, which encode for a matrix protein (Z) and polymerase (L), and a nucleoprotein (NP) and glycoprotein complex (GPC), respectively.^2^ The GPC is the sole protein displayed on the viral surface envelope, and its binding domain, GP1, mediates infection by recognizing the transferrin receptor 1 (TfR1/CD71).^4^ That engagement facilitates internalization into acidified endosomes and, ultimately, viral fusion. Accordingly, the NWM GPC is a focus for antivirals and the GP1, an attractive target for broad neutralization. However, NWM targeting is challenged by the fact that Clade B GP1 sequences vary significantly.

While all pathogenic Clade B NWMs bind human TfR1 (hTfR1), a similarity matrix of GP1 sequences across the clade shows pairwise comparison identities in the 25-50% range (Table S1). Even the closely related Junin virus (JUNV) and Machupo virus (MACV) GP1s are only 48% identical (Table S1), with sequences for the Romero strain (JUNV) and Carvallo strain (MACV) GP1s differing at 101 sites, excluding insertions or deletions; 78 of these lie within the globular portion of the domain that sits atop GP2 (Figure 1). That level of variation impacts the predictability of novel pathogenic variants or emerging zoonoses, since receptor engagement can presumably be accommodated by a large number of GP1 variants.

**Figure 1.**
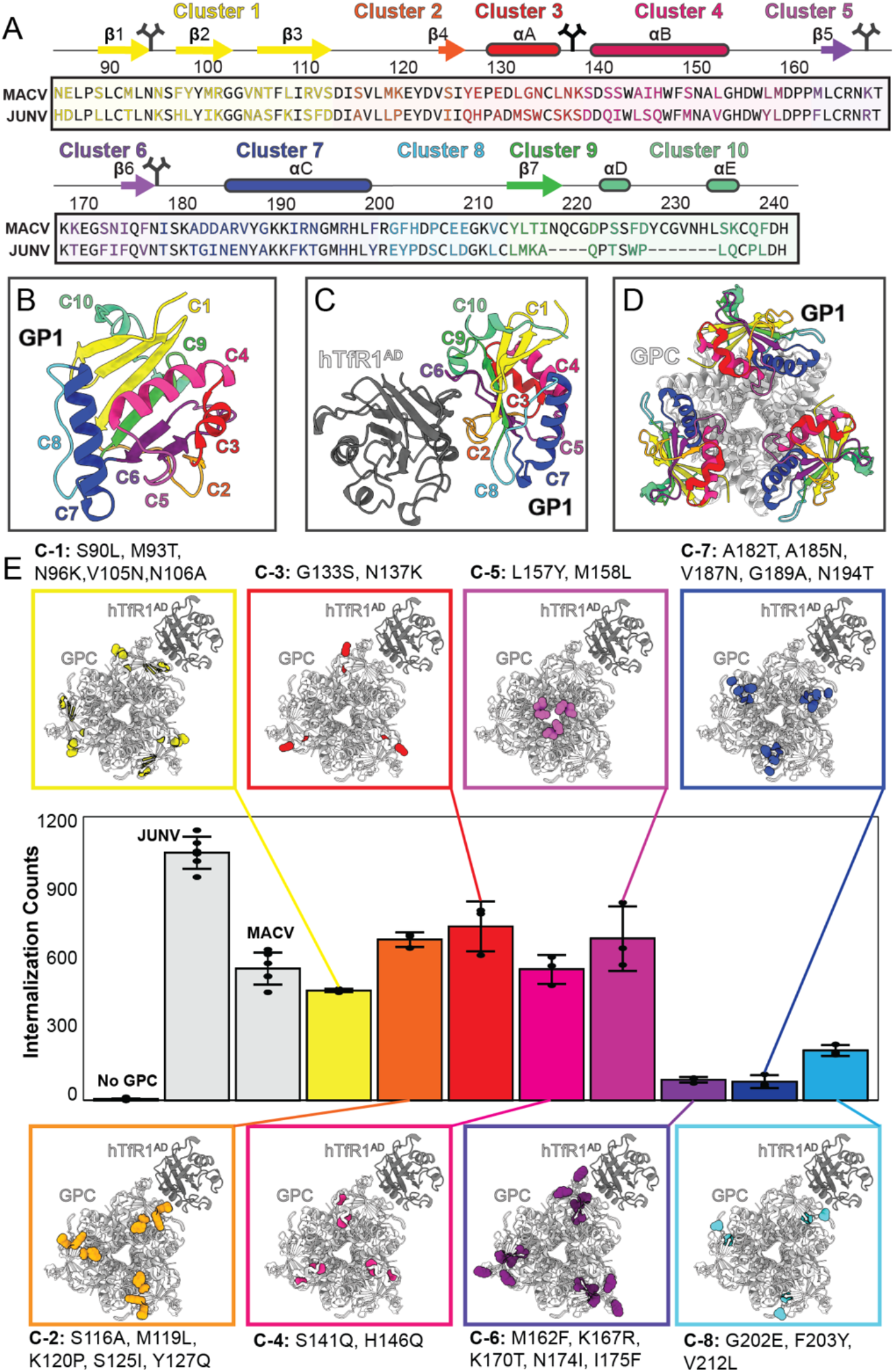
Computational selection of MACV-JUNV GP1 hybrid variants. (A) Sequences of wild type MACV and JUNV GP1 globular domains, with conserved residues colored black, variable residues colored according to one of 10 clusters they reside in. Each cluster is distinctly colored along with its depicted secondary structure elements; glycosylation sites are depicted by branched lines. The structure of MACV GP1 (B), the hTfR1^AD^-bound GP1 (C), and GP1 in the context of the MACV GPC structure (D) are shown with subregions colored by cluster, with colors as defined in (A). (E) Cellular internalization of PVs displaying one computationally selected variant for each of clusters 1-8, colored by cluster. Insets show the affected residues in a model of a MACV GPC trimer docked to hTfR1^AD^.

NWM GP1 sequence variation also challenges the development of broadly effective immunogens and the rational development of antibodies capable of neutralizing multiple NWMs (bnAbs). In contrast to human TfR1-targeting broadly neutralizing antibodies^5^, most GP1-targeting antibodies recognize and neutralize a single virus species. For NWMs, notable exceptions include vaccine elicited cross-reactive antibodies that can neutralize both MACV and JUNV.^6^ These examples motivate a need to further understand the impact of GP1 sequence variation on both its function and immune recognition.

Here we investigate GP1 cryptic sequence variants that bridge closely related pathogenic species, informing on the landscape of combinatorial variation that spans 2^78^ distinct hybridized MACV-JUNV sequences. A structure-informed segmentation of the MACV GP1 sequence into 10 sub-regions reduces the scale of that challenge and permits the computer aided selection of putative TfR1 binders. Biochemical and cellular assays identify 62 cryptic variants that facilitate internalization of pseudotyped virus-like particles (PVs) into hTfR1-expressing cells. Recombinant GP1 constructs of 2 divergent variants are both shown to bind hTfR1 and form stable complexes, adopting receptor-bound structures analogous to that of wild-type MACV. A subset of the functional variants is also recognized by the JUNV-MACV cross-reactive monoclonal antibody CR1-07, but to a lesser extent by a JUNV-specific vaccine elicited antibody or by JUNV convalescent patient plasma. Lastly, immunization of mice with replication competent vesicular stomatitis virus (rVSV) displaying each of six distinct variants reveals the potential for such computationally derived, and structurally and functionally validated variants to elicit cross-neutralizing antibodies against MACV and JUNV.

## Results

### A computational strategy to identify functional cryptic MACV-JUNV GP1 variants compatible with receptor binding

We set out to computationally and experimentally investigate the multidimensional landscape of potential sequence variants bridging the closely related pathogenic NWMs, wild type MACV and JUNV. A residue clustering approach helped reduce the scale of the sequence space search. Variable sites within residues 86-241 in the MACV GP1 structure were partitioned based on linear sequence, secondary and tertiary structure, proximity to the hTfR1 binding site and known neutralizing antibody epitopes (Figure 1). This approach led to the construction of 10 clusters: sequence segments with segregated secondary structure elements within the GP1 structure. The central anti-parallel beta sheet in GP1 was split into clusters 1, 9 and 6 (C-1, C-6, C-9), while its surrounding alpha helices were contained in clusters 3, 4, 7 and 10 (C-3, C-4, C-7, C-10); and clusters 2 and 8 (C-2, C-8) included within adjacent loops (Figure 1, Table 1). By this segmentation, hTfR1-proximal residues rested in C-1, C-2, C-6, C-9, and C-10, while C-3, C-4, C-5, and C-7 faced the core of the GPC (Figure 1). Each cluster was independently subjected to computational selection of energetically preferred residues at each of its variable sites. This yielded an initial set of 8 hybridized MACV-JUNV sequences to be functionally assessed, each with a unique combination of MACV/JUNV residues most compatible with a receptor-bound configuration for a given cluster, and its converse. Clusters whose variants did not meet the selection criteria (C9 and C10) were excluded. Individual hybridized GP1 sequences were then incorporated into a MACV GPC construct and assessed for their ability to enable PV internalization into HEK-293T cells (Figure 1).

### hTfR1-mediated PV internalization of hybridized MACV-JUNV GP1 variants

All C1:C8 variants predicted to be compatible with receptor binding allowed internalization of PVs into HEK-293T cells, albeit to varying degrees; all produced higher counts of GFP-positive cells than the no GPC control (Figure 1). This led us to hypothesize that an iterative, structure-informed combinatorial approach could facilitate the broader selection of hybridized sequences that contained variable residues across clusters: starting from a single preferred variant in C2 with a high degree of cellular internalization, and adding variation in preferred sites across C1:C10 (Figure 2, Figure S1). As done previously for single cluster variants, each new variant was subjected to computational prediction of energetically preferred residues at each of its variable sites (Figure S1). This approach showed that the degree of internalization exhibited by a given variant did not necessarily correlate with the number of its residues that differed from the wild-type MACV sequence, or with the number of clusters allowed to vary. It suggested a high degree of roughness in the sequence landscape, when optimizing for receptor-binding and internalization (Figure 2). Most variants that facilitated cell entry did so to an extent greater than 25% that of JUNV and MACV, and included mutations in C2, C3 and C4 (Figure 2, Table S2). The internalization of all variants of interest was judged to be hTfR1-dependent, given its uniform reduction in the presence of the hTfR1-specific monoclonal antibody, OKT9 (Figure S3) and their specific internalization into CHO cells stably expressing hTfR1 (CHO-TfR1) relative to those that express no TfR1 (CHO-Neo) (Figure S3). That shared reliance on hTfR1 was poorly estimated by AlphaFold3, which did not consistently identify the expected interface between hybridized variants and the hTfR1-apical domain (hTfR1^AD^) (Figure S4).

**Figure 2.**
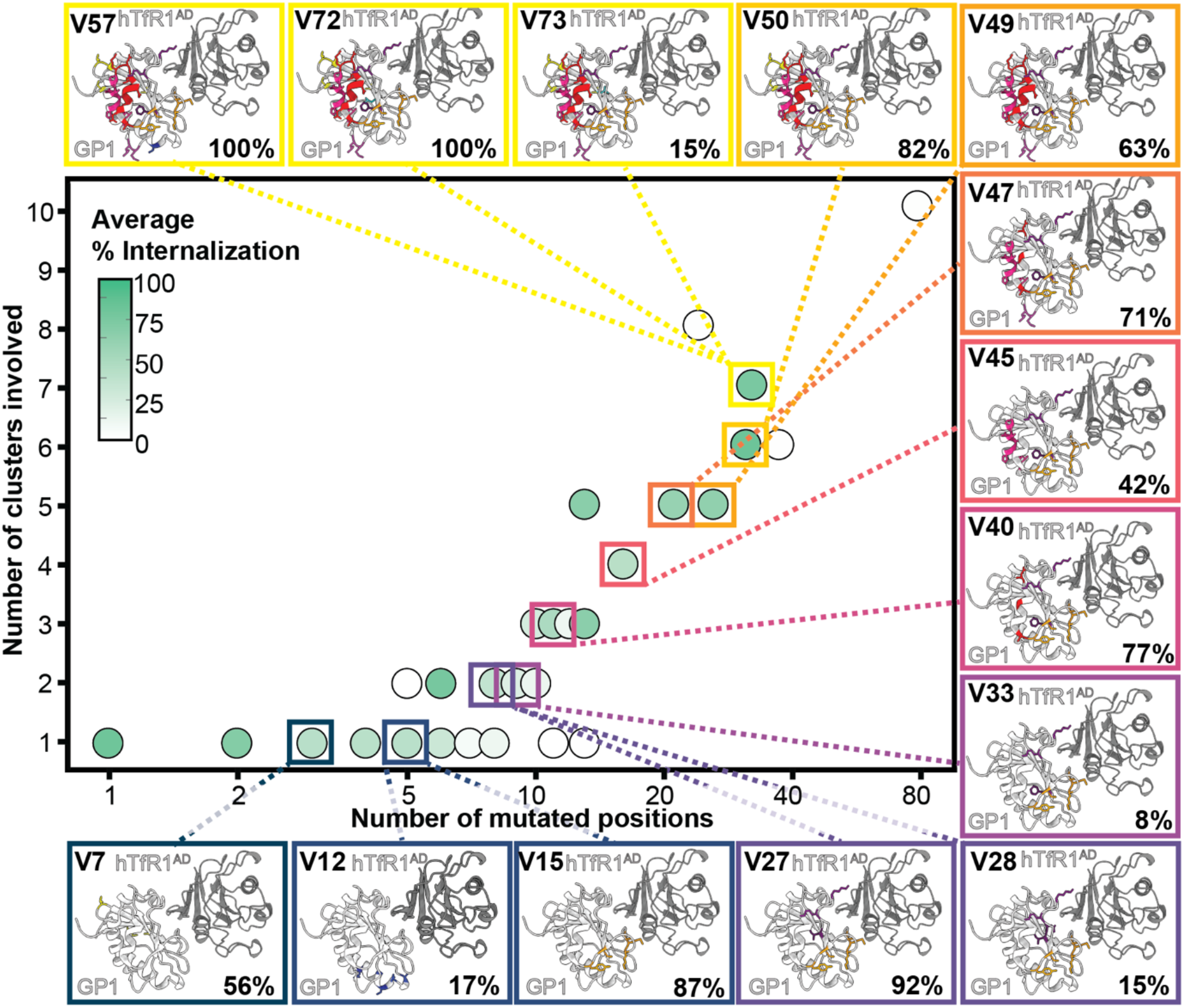
Combinatorial selection of functional GP1 variants across clusters. The number of altered residues per variant is charged against the number of clusters they span in its sequence. Each circle represents a computationally selected variant with its color indicating the level of internalization compared to wild type MACV, from 0% white to 100%, green. Where multiple variants map to the same number of mutations and clusters, the color represents the average level of internalization across each variant at that position. Insets show the altered residues for select variants in the context of the wild type MACV GP1 structure (white) bound to hTfR1^AD^ (gray). Altered residues are colored by their corresponding cluster.

Among the most effective variants was V27, a sequence identified in early iterations that bore 8 variant sites spread across 2 clusters adjacent to the hTfR1-binding epitope, C2 and C6. Also among highly functional variants was V57, a later iteration variant that contained 32 variable sites across 7 clusters (C1:C7) (Table S2). Its existence indicated that bridging local minima in the JUNV/MACV landscape can help identify sequences that match wild-type activity but are significantly hybridized away from either parent wild-type JUNV or MACV sequence (Figure S1, Table S2).

### Recognition of MACV-JUNV GP1 variants by JUNV-selective and JUNV-MACV cross-reactive monoclonal antibodies, and by JUNV reactive convalescent patient plasma

Despite their anticipated structural similarity, we posited that cryptic sequence variants may be engaged by neutralizing antibodies with varying efficiency. We therefore evaluated the recognition of GP1 variants by JUNV-selective antibodies, CR1-28 and convalescent JUNV patient plasma, compared to the JUNV-MACV cross-neutralizer, CR1-07. Given the diversity of variable GP1 positions across the computationally selected set of variants, we assessed the impact of neutralizing antibodies on 14 select variants (Figure 3). Internalization of PVs decorated with each of the 14 variants into HEK-293T cells was generally impacted by both CR1-07 and CR1-28 (Figure 3). However, a dot blot showed all 14 variant PVs were recognized by CR1-07 and most were very weakly recognized by patient plasma; only a subset of variants, including V15, V27, V40, V45, V47, V50, and V57 were recognized by CR1-28 (Figure 3). This indicated that some variants may have unique features that are more robustly recognized by vaccine-elicited cross-neutralizing antibodies, and that even relatively weak recognition by CR1-28 could impact PV internalization. The discrepancy in recognition by the neutralizing antibodies was well exemplified by their interaction with V27 and V57. Recombinant constructs of these variants were bound differentially, in a concentration-dependent manner, by CR1-07 and CR1-28 (Figure 3). That difference may be a consequence of varying residues near the CR1-28 binding site, as suggested by computationally predicted models of the antibodies in complex with the variant GP1s (Figure 3). Those models indicate that recognition by CR1-28 may require remodeling of residues proximal to the C-terminal insertion loop unique to the MACV GP1, where a steric clash between loop residues and CR1-28 may be alleviated by mutations near their interface to allow binding.

**Figure 3.**
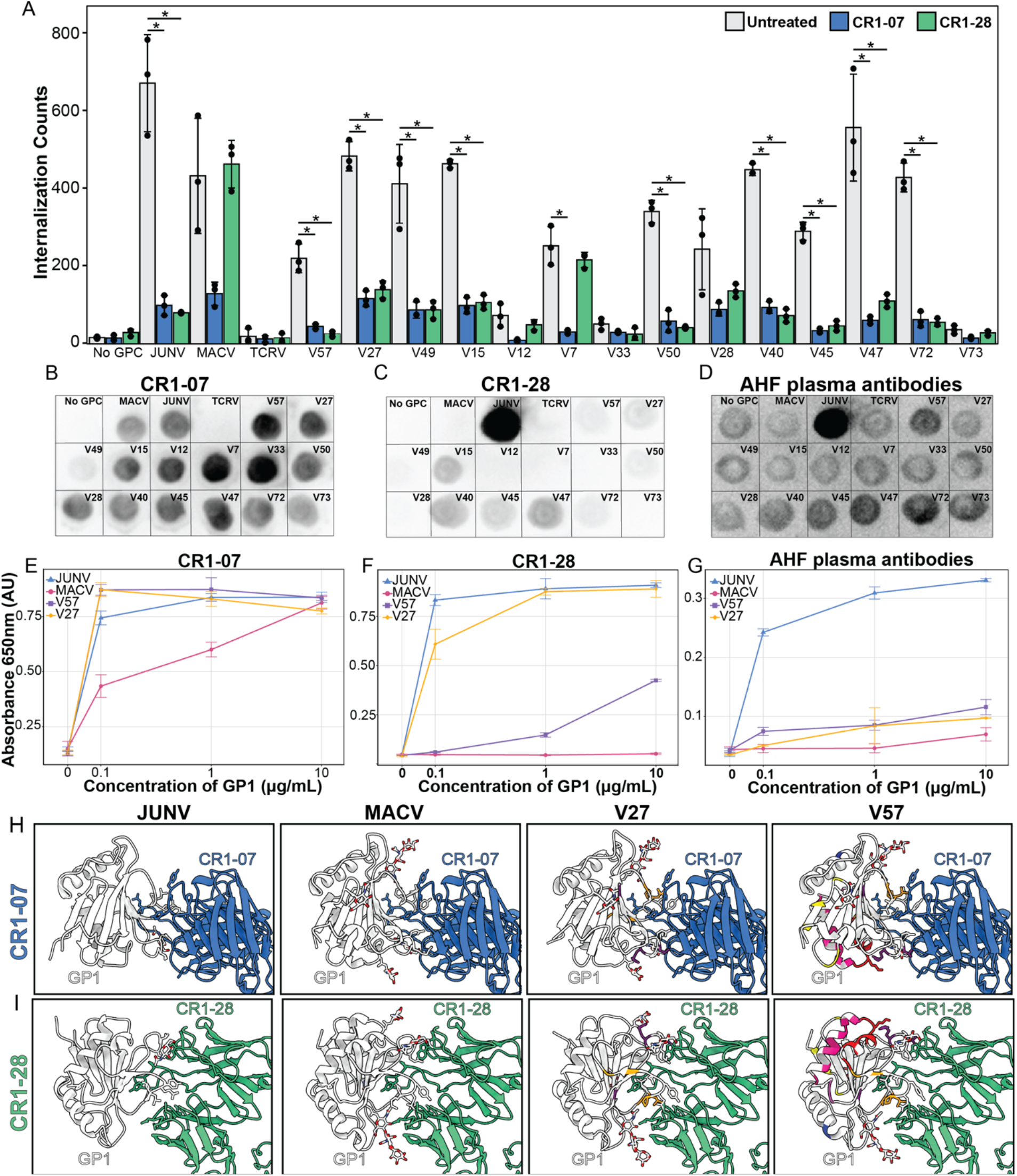
Interactions of cryptic GP1 variants with NWM-neutralizing monoclonal antibodies and recognition of cryptic GP1 variants by anti-JUNV patient sera. (A) Cellular PV internalization assay displaying select GPC sequence variants in the presence and absence of 50µg/mL CR1-07 and CR1-28 monoclonal antibodies. *=p-value <0.05. (B-D) Dot blots of PV media from panel A against CR1-07 (B), CR1-28 (C), and anti-JUNV patient plasma polyclonal antibodies (D). (E-G) ELISA binding assays of recombinantly generated WT and mutant GP1s against CR1-07 (E), CR1-28 (F), and anti-JUNV patient plasma polyclonal antibodies (G). (H-I) Structural models of WT and mutant GP1s in complex with CR1-07 (H) and CR1-28 (I) through alignment with PDBs: 5W1K, 5W1M.^6^

### Structural basis for hTfR1 engagement by computationally hybridized MACV-JUNV GP1 variants

To better understand the structural basis for function and neutralization of V27 and V57, we sought molecular structures of their recombinant soluble forms (GP1^V27^ and GP1^V57^) bound to the soluble ectodomain of hTfR1 (Figure S2). We prepared complexes of holo human transferrin (hTf)-hTfR1 bound to recombinant wild type MACV GP1, GP1^V27^, or GP1^V57^ (Figure S2) in a 1:3 stoichiometry. These appeared monodisperse by negative stain EM and suitable for cryoEM analysis (Figure S6). Single particle reconstructions revealed wild-type MACV and V27 GP1-hTfR1 complexes to be C2 symmetric (Table S3), with partial occupancy for their GP1 subunits (Figure S7-8), while V57 showed partial occupancy for only one lobe of hTfR1 and was refined using C1 symmetry (Table S3, Figure S9). In the absence of complete and high-resolution density for V57, the model was rigid body fit and real space refined into the cryoEM map. Although V57 and V27 both differ by four receptor-proximal residues from wild-type MACV GP1 (S116A, M119L, K120P, K170T), they and the wild type construct bound hTfR1 in nearly identical configurations (Figure 4). The variants showed overall RMSDs of ∼0.5 Å with respect to wild type, with each receptor-proximal altered residue being accommodated by its corresponding structure (Figure 4). What was not apparent from the determined structures was the impact of the extensively hybridized residues at the GP1-GPC interfaces on the overall structure and integrity of the full length GPC itself, although the soluble variant constructs were no less stable than their wild type counterpart (Figure S2), and both variants can form functional GPCs capable of internalizing through hTfR1 (Figure S3).

**Figure 4.**
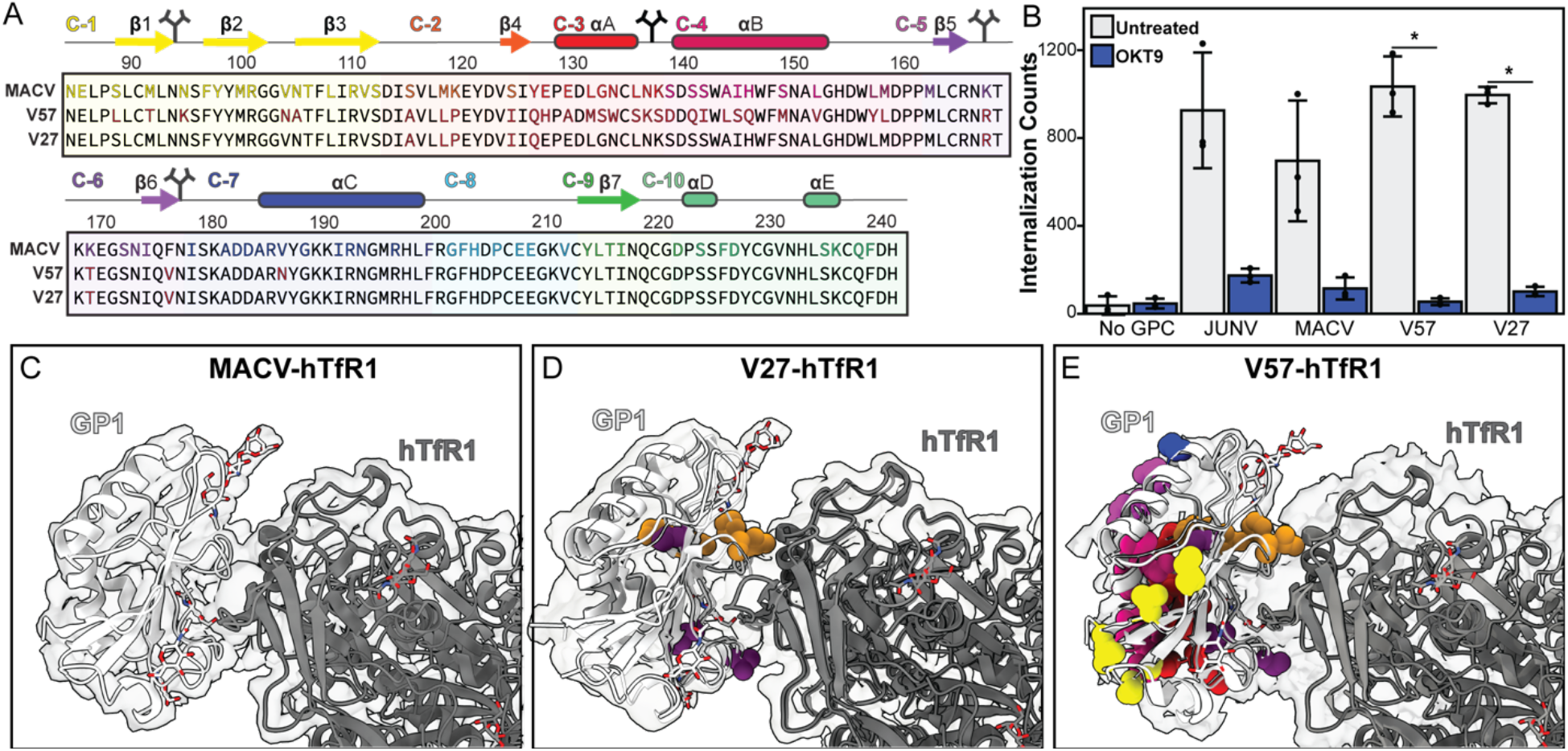
Structural and functional characterization of two hybridized JUNV-MACV GP1 cryptic variants. (A) Sequences of two cryptic GP1 variants alongside the wild-type sequence of the MACV GP1 globular domain. Conserved residues are colored black; in the wild-type sequence, variable residues are colored based on their cluster, while in the variant sequences, they are colored brick red. (B) Cellular internalization of PVs displaying no GPC, wild-type JUNV, MACV GPCs, or the MACV GPC with altered residues according to each of the two variants. Each condition is shown with and without incubation with the hTfR1-targeting antibody, OKT9. *=p-value <0.01. (C-E) Single particle cryoEM structure of wild type soluble MACV GP1 construct (C) V27 GP1 construct (D), or V57 GP1 construct (E) bound to hTfR1; altered residues are shown as space fill models, colored by their corresponding cluster.

### Eliciting cross-reactive neutralizing antibodies with variant GP1-bearing GPCs as designed immunogens

The functional engagement of variants with hTfR1 and their recognition by cross-neutralizing antibodies raised their potential as immunogens to elicit JUNV-MACV cross-reactive neutralizing antibodies. To assess this, mice were immunized with VSV PVs displaying MACV-Carvallo, JUNV-Romero, or variant GP1-bearing GPCs. Immunizations generated robust humoral responses as indicated by ELISA-based antibody titers against the targets, with V12, V15 and V27 inducing pronounced responses against both wild type JUNV and MACV GPCs (Figure 5). In contrast, V49 and V57 elicited antibody responses that showed weak titers against either JUNV GPC, MACV GPC or GP1. Overall, while cross reactive titers could be elicited by several variants, and weak inhibition of VSV PVs bearing JUNV-Romero was observed for all variants, MACV-Carvallo PVs was only inhibited using sera elicited by four variants: V7, V12, V15 and V27 (Figure 5). The level of detection of MACV or JUNV GPC by sera showed a variable association with partial protection against JUNV or MACV PVs, with V27-elicited sera showing moderate cross-reactive inhibition of JUNV and MACV PVs, while V57 showed only moderate inhibition of JUNV PVs (Figure 5). This selectivity indicates a relatively higher level of background cross-neutralization induced by MACV vs. JUNV, and the challenge of identifying variants that not only elicit JUNV/MACV cross-reactive antibodies, but also potently cross-neutralize both JUNV and MACV.

**Figure 5.**
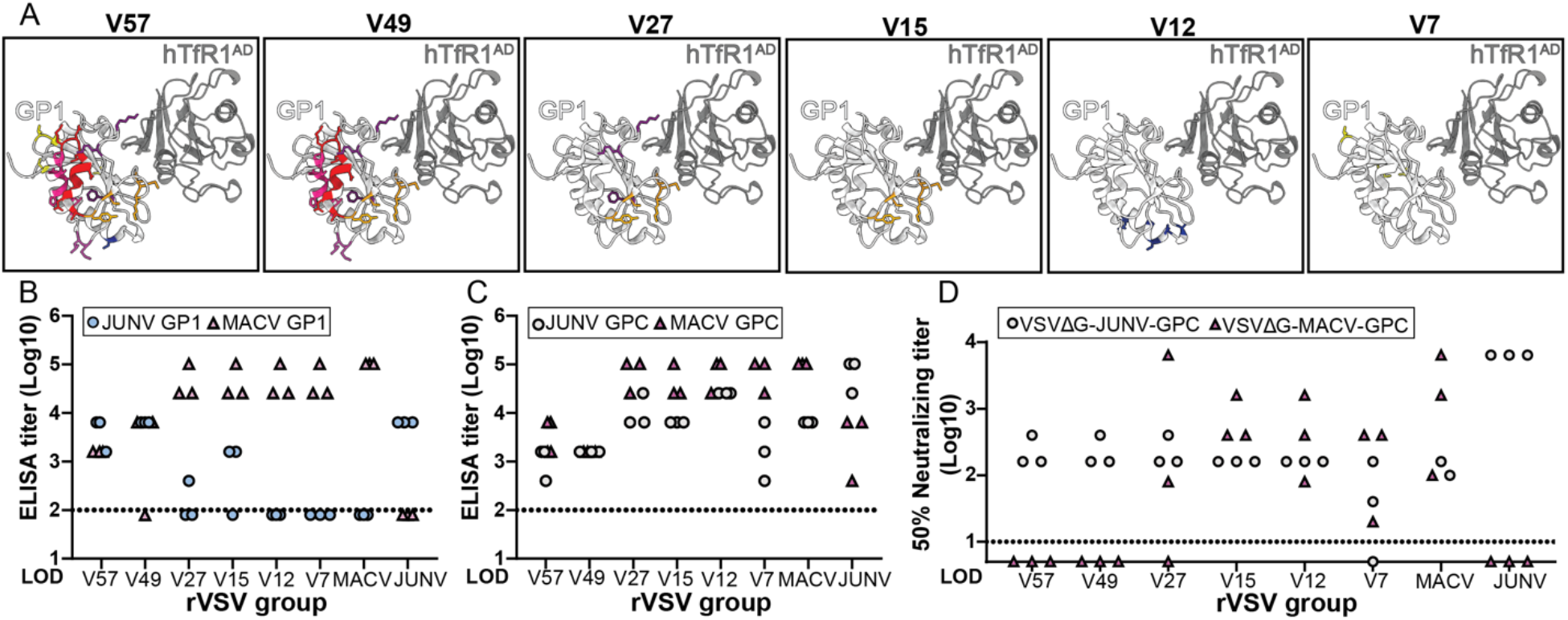
Cross neutralizing response of cryptic GP1 variants produced through *in vivo* immunization. (A) Models of select GP1 variants (white) in complex with hTfR1^AD^ (grey) with mutations relative to MACV colored by their corresponding cluster used for *in vivo* immunization experiments. (B-C) ELISA binding titers of unpurified polyclonal serum antibodies collected 2 weeks post-infection derived from WT (MACV and JUNV) and mutant GP1-VSV pseudotyped virus infections to recombinantly generated MACV and JUNV isolated GP1 domains (B) and GPC trimers (C).The endpoint titer was defined as the highest dilution that exceeded an OD450 value of 0.1. (D) Neutralizing activity of polyclonal antibodies in the serum samples collected 2 weeks post-infection, calculated by determining the sera dilution that resulted in a 50% reduction in infectivity of PV compared to a control well without sera.

## Discussion

The sequence space separating the GP1 subunits of closely related pathogenic JUNV and MACV species is remarkably vast, despite their similar structures and functions. Out of ∼2^78^ potential GP1 variants, we anticipated that a subset might retain their native structure, rely on hTfR1 for cellular entry, and elicit cross-neutralizing immune responses, making them promising vaccine candidates. To characterize cryptic JUNV/MACV hybrid GP1 variants, we relied on their computational prediction and biochemical and cellular validation of function. This approach revealed a rugged landscape with function not generally predicated on the number or location of varying sites on GP1. Instead, an iterative approach uncovered a path to functional variants with an increasing number of altered sites away from the wild-type MACV sequence, bearing as many as 32 JUNV residues. A subset of these variants was recognized by the JUNV/MACV cross-neutralizing antibody CR1-07, and surprisingly, several were also bound by the JUNV-specific antibody CR1-28. Models suggest the structural basis for this gained recognition is in part due to interactions proximal to a C-terminal insertion loop unique to the MACV GP1. Despite their distinct pattern of engagement with neutralizing antibodies, two of these variants, V27 and V57, were also found to closely resemble wild-type MACV in their binding to hTfR1. Some hybridized variants grafted into PV-based immunogens for mouse challenge were capable of eliciting cross-reactive and cross-neutralizing antibody responses. Chief among these was GP1^V27^, which differed from wild-type MACV GP1 at 8 sites, but showed promise in eliciting both cross-reactive and cross-neutralizing antibodies; variants with more altered sites lost the ability to elicit MACV neutralizing antibodies.

Recognition of functional JUNV/MACV hybrid GP1 variants by cross-neutralizing antibodies, such as the vaccine-elicited CR1-07 or those in the sera of immunized mice was not generally predictive of the neutralizing potential of the antibodies against MACV PVs. This is exemplified by the sera of mice vaccinated with V49 and V57, which recognized JUNV and MACV GP1 or GPCs by ELISA, but were unable to neutralize MACV PVs in culture. This could be due to the high number of JUNV residue-containing sites present in these variants, 26 for V49 and 32 for V57 (Table S2). Those differences may have steered the responses toward epitopes that could be targeted to preclude receptor engagement by JUNV but not MACV. GP1^V27^ stands in contrast to these, with fewer altered sites and the capacity to elicit some cross neutralizing antibodies. A benefit of the iterative computational hybridization approach is the ability to identify variants that represent disparate islands of stability in the sequence-function landscape, driving cross-reactive responses. The present strategy achieves this by hopping between these islands of stability, but lacks the more exhaustive, dense mutational analyses that might rationalize their existence.^7–10^

Deep mutational scanning (DMS) has been highly successful in comprehensively mapping single and variation and expanding into multi-site variation in other viral glycoproteins.^7–10^ These have yielded models informed by a richness of sequence-function links. Such efforts on NWM GPCs would complement the current approach and potentially give rise to a more nuanced selection of GP1 variants that elicit cross-neutralizing immune responses. While we observed such responses from computationally selected immunogens, their polyclonal nature obfuscated prediction of cross-neutralization potential. As computational models improve, those trained specifically on the behavior of variants as immunogens could yield better predictions and permit the more routine design of structure-based vaccines.

## Acknowledgments

We thank Drs. Michael Sawaya and Duilio Cascio (UCLA) for assistance, and S2C2 personnel, Alexandre Cassago, Ian Fries, for their invaluable support and assistance. We also thank Drs. Silvana C. Levis, Ana M. Briggiler and Delia A. Enria for providing Argentine hemorrhagic fever survivor plasma samples from Instituto Nacional de Enfermedades Virales Humanas “Dr. Julio I. Maiztegui”, (INEVH, Pergamino, AR). This work was supported by NIH-NIGMS Grant R35GM128867 to J.A.R., the Howard Hughes Medical Institute Emerging Pathogens Initiative (HHMI EPI), and was also performed as part of STROBE, an NSF Science and Technology Center through Grant DMR-1548924. J.A.R. was further supported as a Packard Fellow. Single particle cryoEM was acquired at the Electron Imaging Center for Nanosystems (EICN) at the UCLA California NanoSystems Institute (CNSI) (RRID:SCR_022900) (NIH S10OD032459), and at the Stanford-SLAC Cryo-EM Center (S2C2), which is supported by the National Institute of General Medical Sciences (1R24GM154186). J.M. was supported by internal funds from the Department of Pathology at UTMB, the Institute for Human Infections and Immunity at UTMB, the John S. Dunn Foundation, and NIH R00AI156012. G.S. was supported by the Long Term Health and Education Training (LTHET) Program funded by the U.S. Army. The views and information presented are those of the authors and do not represent the official position of the U.S. Army Medical Center of Excellence, the U.S. Army Training and Doctrine Command, the Department of the Army, the Department of Defense, or the U.S. Government.

## Data availability

All data and protocols related to this study will be made available without restriction by the authors. Models and maps of hTfR1 in complex with wild-type MACV GP1, V27 GP1, or V57 GP1 are deposited under PDB accession codes 12HO (wild-type MACV GP1), 12HS (V27), 12HT (V57) and EMDB entries EMD-76440 (wild-type MACV GP1), EMD-76443 (V27), EMD-76-444 (V57).

## Author contributions

J.A.R. and L.J.T. conceptualized and directed the work. L.J.T. computationally generated and validated sequence variants. L.J.T., S.F., and G. S-T. performed biochemical and cellular experiments. L.J.T. prepared, collected, and analyzed cryoEM data, and performed structure determination. S.F. and G.H. procured and prepared convalescent patient plasma. T.S. and G.S-T. prepared rVSV particles. G.S-T. and J.M. conducted VSV immunization and associated mouse work. L.J.T., and J.A.R. wrote the article, with input from all authors.

## Materials & Methods

*Materials and reagents*

### Ethics statement

All animal studies were reviewed and approved by the Institutional Animal Care and Use Committee (IACUC) at UTMB and conducted in accordance with the National Institutes of Health guidelines. Experimental procedures involving replication-competent vesicular stomatitis Indiana virus (rVSV) were conducted within the Animal Biosafety Level 2 (ABSL-2) facility at UTMB. All procedures were performed in accordance with institutional ethical guidelines and the AVMA Guidelines for the Euthanasia of Animals (2020 Edition), ensuring animals were fully unconscious and insensible prior to death. All efforts were made to minimize animal suffering.

### Cells

HEK-293T/293TT (human embryonic kidney epithelial) cells were grown in 4:1 OptiMEM to DMEM mix (Life Technologies; Thermo Fisher Scientific) supplemented with 1-2% fetal bovine serum (FBS, Life Technologies; Thermo Fisher Scientific) and 1% penicillin-streptomycin (100 U/ml penicillin and 0.1 mg/mL streptomycin) in a humidified incubator at 37°C with 5% CO_2_.

CHO-Neo and CHO-TfR1 cells^11^ were grown in DMEM (Life Technologies; Thermo Fisher Scientific) supplemented with 10% fetal bovine serum (FBS, Life Technologies; Thermo Fisher Scientific) and 1% penicillin-streptomycin (100 U/ml penicillin and 0.1 mg/mL streptomycin) in a humidified incubator at 37°C with 5% CO_2_.

OKT9 hybridoma cells (ATCC-CRL-8021) were grown in Hybridoma Serum-Free medium (SFM) supplemented with 1% penicillin-streptomycin (100 U/ml penicillin and 0.1 mg/mL streptomycin) in a humidified incubator at 37°C with 5% CO_2_.

African green monkey kidney Vero, were grown in Dulbecco’s modified Eagle’s medium (DMEM) (Corning) supplemented with 10% fetal bovine serum (FBS) (Avantor), 100 U/mL penicillin, and 0.1 mg/mL streptomycin (Gibco). These cells were cultured at 37°C with 5% CO_2_. Baby hamster kidney BSR-T7/5 cells^12^ were maintained in DMEM containing 1 mg/mL of geneticin and 10% FBS. The BSR-T7/5 cells were passaged to avoid reaching over 80% confluency. Expi293F cells were grown in Expi293™ expression medium (Gibco) at 37°C with 8% CO_2_, 80% humidity, and on an orbital shaker platform at a constant rotation of 125rpm.

### Plasmids

MACV-JUNV GPC sequence variants were generated through cloning of mutated residues into the MACV GPC codon-optimized sequence from the Carvallo virus strain within the mammalian expression vector pcDNA 3.1+ by Twist Bioscience. PVs displaying WT and MACV-JUNV sequence variants were generated as previously described^13–15^. Briefly, each GPC plasmid was co-expressed alongside the MLV gag pol polyprotein and a packaging-competent eGFP appended to an N-terminal SV40 nuclear localization signal in 293TT cells. Full plasmid sequences were obtained for all constructs through Plasmidsaurus sequencing.

Viral cDNA encoding the GPC gene from the MACV strain Carvallo and the JUNV strain Romero was generated from extracted viral RNA by reverse transcription using SuperScript IV Reverse Transcriptase (ThermoFisher) with gene-specific primers according to the manufacturer’s instructions. The PCR-amplified MACV, JUNV, or mutant GPC genes were inserted into the pVSV-ΔG-PL 2.5 vector (Kerafast) using the In-Fusion cloning kit (Takara Bio) according to the manufacturer’s instructions. The expression plasmids pBS-VSV-N, pBS-VSV-P, and pBS-VSV-L were purchased from Kerafast.

The pCAGGS expression plasmids for recombinant soluble MACV or JUNV-GPC (pCAGGS-MACV GPCTri and pCAGGS-JUNV GPCtri) were constructed as previously^16^. Briefly, each gene was amplified with KOD One PCR Master Mix (Takara Bio) and gene-specific primers, and cloned into pCAGGS using an In-Fusion Snap Cloning kit (Takara Bio) according to the manufacturer’s instructions. The sequence of all the plasmids were confirmed by Next Generation Sequencing (Genewiz). The expression plasmid for His-tagged Ebola virus GP^17^ was kindly provided by Dr. Ayato Takada, Hokkaido University International Institute for Zoonosis Control.

### Prediction of functional MACV/JUNV sequence variants

Using a fast-relaxed, trimeric model of the hTfR1^AD^-MACV GPC complex, residues corresponding to Junin and Machupo residues 86-241 were computationally sampled at each non-conserved site within each cluster region using Rosetta design.^18^ Generated sequence variants, holding mutations within each cluster regions, were scored and selected on the basis of total complex score and interface score (sum of per residue energies for each binding residue at the hTfR1 interface) using the Rosetta energy function. Sequence variants with predicted function were selected for experimental validation alongside inverse variants where all mutations selected through energetic favorability were maintained as the WT MACV and mutations not selected were converted to the corresponding JUNV residue. The relative internalization of each sequence variant was assessed as a percentage of internalization relative to WT MACV through quantification of GFP expression. After experimental validation using pseudovirus internalization assays, this process was iterated as mutations were fixed in one cluster region at a time.

### Pseudotyped virus-like particles production for internalization assays

Pseudotyped virus-like particles internalization assays were conducted as previously described.^15^ Briefly, 293TT cells were plated at 70-80% confluence in a 96-well plate then transfected with MLV gag/pol, GFP, and GPC plasmids using Fugene transfection reagent. After washing the cells and incubating for 60-72 hours, PV-containing media was transferred directly from the expressing cells to fresh HEK293T cells at 50% confluence in a 96-well plate. The cells were washed 16 hours post addition of PV media and NucBlue live cell straining solution was added. Images of the cells were collected at 20X magnification using an Evos M7000 wide-field fluorescence light microscope for both DAPI and GFP channels covering 12-20% of the surface area of each well. Images were automatically stitched together into montages by the Evos analysis software and unique GFP events were quantified using a custom image analysis MATLAB script.

### Production of antibodies

#### CR1-28 and CR1-07

Plasmids encoding the heavy and light chains of CR1-28 and CR1-07^19^ antibodies were provided by Jonathan Abraham and were prepared as previously described.^15^ Briefly, heavy and light chain plasmids of CR1-07 and CR1-28 were expressed in HEK293T cells. After one week, the cell culture media was collected, clarified by centrifugation, and filtered. The antibodies were purified from the filtered media using a protein G column and eluted with a Glycine buffer at pH 2.7 Purified fractions were dialyzed into TBS and concentrated to approximately 2mg/mL.

#### OKT9

OKT9 was generated as previously described.^15^ Briefly, Hybridoma cells stably expressing OKT9 were grown in SFM for one week. The cell culture media was then collected, clarified by centrifugation, and filtered. OKT9 was purified from the filtered media using a protein G column and eluted with a Glycine buffer at pH 2.7 Purified fractions were dialyzed into TBS and concentrated to approximately 5mg/mL.

#### Polyclonal AHF plasma-derived antibodies

Total IgG was purified from human plasma samples of AHF convalescent patients using affinity chromatography. Briefly, high-titer plasma (1:10,240) was clarified by centrifugation and filtered through a 0.45 µm membrane. The sample was then diluted 1:4 in binding buffer (TBS: 50 mM Tris, 137 mM NaCl, 2.7 mM KCl, pH 7.6). Affinity purification was performed using ProteinIso® Protein G Resin (TransGen Biotech). The resin was equilibrated with TBS, and the diluted plasma was incubated to allow IgG binding. After washing the resin with TBS, the bound IgGs were eluted using elution buffer (0.1 M glycine-HCl, pH 2.5). Each fraction was immediately neutralized by adding 1 M Tris-HCl pH 9.0. All separation steps, including resin equilibration, washing, and elution were performed by centrifugation at 1,000 xg for 1 minute to collect the supernatants. Protein concentrations in all fractions were determined using a NanoDrop spectrophotometer.

#### cryoEM Grid Preparation

2.5 µL of WT or mutant GP1 in a 3:1 stiochiometric ratio with sTfR1 were applied to glow-discharged (PELCO easiGLow -Ted Pella) R2/1 Quantifoil grids and placed in a Vitrobot Mark IV (Thermo Fisher Scientific) at 10°C and 100% relative humidity and blotted using a blot force of 0 and blot time of 0.5-1.5 seconds, and plunge frozen into liquid ethane. cryoEM grids were screened for optimal ice and particle distribution using a TALOS F200C TEM (Thermo Fisher Scientific) operated at 200 KeV and equipped with a direct electron detector (Apollo – Direct Electron). cryoEM grids with favorable ice conditions were clipped using a SubAngstrom clipping station and stored in liquid nitrogen for high-resolution imaging.

#### cryoEM Data Acquisition

cryoEM data acquisition of GP1-sTfR1 complexes was performed on the UCLA CNSI Titan Krios (MACV, V57) or the Stanford SLAC Titan Krios (JUNV, V27) operated at 300 KeV using a Falcon4i electron detector. Movies were collected in super resolution mode with a calibrated pixel size of 1.2 Å (MACV, JUNV, V57) or 0.95 Å (V27) per pixel, over a target defocus range of -0.5 to -1.5 µm and a total dose of ∼40 e^-^/ Å ^2^. Movies across datasets were recorded with EPU software Thermo-Fisher Scientific. ^11^

#### cryoEM Data Processing

CTF estimation was conducted on motion-corrected micrographs. Micrographs were curated for further analysis. An initial set of particles for each dataset was picked through blob picker, subclassified, and used to train a Topaz model. Particles were automatically picked using the trained models. 2D classification was performed followed by ab initio reconstructions. The resulting reconstructions served as the reference for 3D classification and refinements without applied symmetry. Next, homogenous and non-uniform refinements were performed, enforcing C2 symmetry for MACV and V27-bound structures, while the V57 dataset was refined using C1. Additional 3D classification without alignment was used to identify the best-resolved classes with the highest GP1 occupancy, which were subjected to additional local refinements using a mask that included the apical domain of hTfR1 and the bound GP1, as well as global and local CTF refinements. Particles were then subjected to reference-based motion correction using the 3D reconstructions in cryoSPARC and further refined with non-uniform refinement with applied C2 (MACV, V27), or C1 (V57) symmetry. An overall resolution of 2.5 Å for both MACV and V27, and 2.7 Å for V57, structures was achieved for the entire particle, based on the 0.143 FSC threshold criterion. Local refinement on the apical domain of hTfR1 and the bound GP1 yielded resolutions of 2.4 Å for both MACV, 2.6 Å for V27, and 3.3 Å for V57, based on the 0.143 FSC threshold criterion.

#### Model Building

An initial model of each GP1-hTfR1 complex was of the was generated by docking the MACV GP1 (PDB: 3KAS)^20^ into a model of hTfR1 with bound transferrin (PDB: 11ZB). The initial models were docked into the cryoEM density in Chimera and further relaxed into the density using Rosetta. Mutations for sequence variants were manually added in Coot. The final model for each complex was built in Coot, then iteratively refined in Phenix. The final models were evaluated for proper geometric properties in Coot and for fit to the density by Q-score.

#### Dot Blots

Dot blots were performed as previously described.^15^ Briefly, PDVF membrane was activated in methanol and equilibrated in TBS-T buffer. PV-containing media or purified protein was added to the membrane and incubated prior to the addition of blocking buffer (5% NFDM in TBS-T). The membrane was then incubated with a 1:1000 dilution of primary antibodies at 1-2 mg/mL for 2 hours. Then membrane was washed in TBS-T then incubated with secondary antibody, and washed in TBS-T. The membrane was incubated in chemiluminescence substrate for 5 minutes then imaged using an Azure biosystems chemiluminescence imager.

#### Viruses

Replication-incompetent pseudotyped vesicular stomatitis Indiana virus (VSV) containing the green fluorescent protein (GFP) gene instead of the VSV-G gene bearing GPCs of MACV strain Romero, or MACV strain Carvallo (VSVΔG-MACV-Carvallo-GPC, and VSVΔG-JUNV-Romero-GPC, respectively) were generated as described previously^21–23^. Briefly, HEK293T cells were transfected with the pCAGGS expression plasmid containing the GPC genes, and 24 hours later, the cells were infected with VSVΔG-VSV-G at a multiplicity of infection of 3.0. After 16 hours of incubation, the supernatants were collected and centrifuged to remove cell debris. Infectious units (IU) of the pseudotyped viruses were determined by inoculating confluent Vero cell monolayers in 96-well plates with 10-fold serially diluted pseudotyped viruses and counting GFP-expressing cells 24 hours later using Opera Phenix Plus System (Revvity). To reduce the background infectivity of the residual parent VSVΔG-VSV-G, each pseudotyped virus stock was treated with a neutralizing monoclonal antibody specific for the G protein (VSV-G[N]1-9)^24^ before use.

The replication-competent vesicular stomatitis Indiana virus (rVSV) expressing different GPCs was rescued from plasmids as described previously with some modifications^25,26^. The BSR-T7/5 cells were seeded on a poly-L-lysine-coated 6-well plate at 1.5 × 10^5^ cells/mL in 2.0 mL of DMEM supplemented with 10% FBS and no antibiotics one day before transfection. The cells were transfected with a mixture of 200 µl of Opti-MEM, 12 µl of FuGENE 4K (Promega), 0.5µg of pBS-VSV-N, 1.25 µg of pBS-VSV-P, 0.25 µg of pBS-VSV-L and 2.0 µg of pVSV encoding the different GPCs. After 6 hours post-transfection, the medium was replaced with DMEM supplemented with 10% FBS and no antibiotics. After 3 days post-transfection, the supernatant was collected and used to inoculate Vero cells. After observation of cytopathic effect, viral RNA was purified from the cell culture supernatant using the Viral RNA Mini Kit (Qiagen), followed by PCR amplification. Each GPC gene sequence was confirmed by Sanger sequencing (Genewiz) and found to have no mutations. The virus was further propagated in Vero cells and stored at −80°C until use. Virus titers were determined by a plaque assay. Confluent Vero cells in 12-well plates were infected with 10-fold serial dilutions of virus stocks and incubated for 30 minutes at 37°C with 5% CO_2_. After the inoculum was removed, EMEM containing 2% FBS, 1% penicillin-streptomycin, and 0.6% tragacanth (Sigma) was overlaid. After 3 days of incubation, cells were fixed with 10% formalin and stained with 0.25% crystal violet. Viral titers were represented as PFU.

#### *In vivo* immunization

Five-week-old female BALB/c mice were purchased from Charles River. All animals were housed in animal biosafety level-2 (ABSL-2) facilities in the Galveston National Lab (GNL) at UTMB. The mice in each group (n=3) were infected twice intraperitoneally (i.p.) with up to 10^6^ PFU of their respective rVSV three weeks apart. Two weeks after the second infection, sera was collected and used in neutralization tests and ELISA-based binding assays.

### Enzyme-linked immunosorbent assay (ELISA)

#### GP1-antibody ELISA binding assays

1-5 µg/mL purified antibody was added to immunogen 96 well plates in coating buffer (carbonate buffer pH 9.0) overnight at 4°C. The plate was then washed 3 times with PBS-T and then incubated with blocking buffer (PBS-T + 1% BSA + 5% NFDM) for 2 hours at room temperature. The 96 well plate was washed again 3 times in PBS-T and then incubated with GP1 recombinant protein at room temperature for 2 hours. After washing again with PBS-T 3 times, the plate was then incubated with secondary anti-His HRP antibody at a 1:1000 dilution for 1 hour at 37 °C. The plate was then washed 4 times in TBS-T and incubated with 3,3’, 5,5’-tetramethylbenzidine (TMB) substrate for 20 minutes prior to absorbance reading at 650 nm using a Varioskan plate reader.

#### Immunization serum ELISA binding assays

GPC-based ELISA was performed as described previously^17,21^. Briefly, to detect anti-GPC antibodies, ELISA plates (Thermo Fisher Scientific) were coated with purified recombinant MACV GP1, JUNV GP1, mutant 1 GP1, mutant 3 GP1, MACV, JUNV GPC, or EBOV GP (50 ng/50 µL/well) in PBS at 4°C overnight and then washed with PBS-T. The plates were blocked with 3% skim milk (200 μL/well) for 1 hour at room temperature. After washing with PBS-T, serially diluted sera in PBST containing 1% skim milk were added (50 µL/well) and incubated at room temperature for 1 hour. Plates were washed with PBS-T 3 times, then 1:10,000-diluted HRP-conjugated goat anti-mouse IgG (Invitrogen) was added and incubated for 1 hour at room temperature. After 5 times washing with PBS-T, the reaction was visualized by adding TMB liquid substrate, Supersensitive for ELISA (Sigma). The reaction was stopped by adding 1 N phosphoric acid, and the optical density at 450 nm (OD450) was measured using a Biotek ELx808 Microplate Reader (Agilent) and normalized to the OD450 of negative control antigens. The endpoint titer was defined as the highest dilution that exceeded an OD450 of 0.1.

#### Neutralization tests

VSVΔG-MACV-Carvallo-GPC and VSVΔG-JUNV-Romero-GPC were appropriately diluted to yield 1,000 to 2,000 IU/well and mixed with serially diluted sera for 1 hour at 37°C and inoculated into confluent Vero cells grown in 96-well plates. Twenty hours post inoculation, GFP-positive cells were counted. The relative percentage of infectivity was calculated by setting the number of cells infected in the absence of sera to 100%. The endpoint titer was defined as the highest dilution at which the viral level was 50% lower than in the control well.

