## Supplemental tables and figures for "Computationally hybridized pathogenic mammarenavirus receptor binding domains reveal cryptic variants that elicit cross-neutralizing immune responses"

### Supplementary Tables

**Table S1**. Percent identity matrix of Clade B NWM GP1s

| | TCRV | MACV | JUNV | SABV | CHAV | AMAV | GTOV | CPXV |
|---|---|---|---|---|---|---|---|---|
| **CPXV** | 24.2 | 26.8 | 26.2 | 30.9 | 29.8 | 52.9 | 52.9 | 100 |
| **GTOV** | 24.3 | 30.8 | 30.2 | 27.8 | 31 | 50.8 | 100 | 52.9 |
| **AMAV** | 30.2 | 31.1 | 32.2 | 28.2 | 26.6 | 100 | 50.8 | 52.1 |
| **CHAV** | 23.1 | 29.5 | 28.4 | 58.9 | 100 | 26.6 | 31 | 29.8 |
| **SABV** | 20.3 | 27.3 | 25.1 | 100 | 58.9 | 28.2 | 27.8 | 30.9 |
| **JUNV** | 45 | 47.7 | 100 | 25.1 | 28.4 | 32.2 | 30.2 | 26.2 |
| **MACV** | 44.5 | 100 | 47.7 | 27.3 | 29.5 | 31.2 | 30.8 | 26.8 |
| **TCRV** | 100 | 44.5 | 45 | 20.3 | 23.1 | 30.2 | 24.3 | 24.2 |

% Identity
100
80
60
40

**Table S2**. List of sequence variants and their mutations relative to the MACV GP1

| Variant Number | Mutations relative to MACV GP1 | Clusters involved |
|---|---|---|
| 1 | Y127Q | Cluster2 |
| 2 | G133S,N137K | Cluster3 |
| 3 | S141Q,H146Q | Cluster4 |
| 4 | L157Y,M158L | Cluster5 |
| 5 | S173F,F177V | Cluster6 |
| 6 | M162F,K170T | Cluster6 |
| 7 | M93T,N96K,V105N | Cluster1 |
| 8 | G202E,F203Y,V212L | Cluster8 |
| 9 | S116A,M119L,K120P,S125I | Cluster2 |
| 10 | H204P,P206S,E208L,E209D | Cluster8 |
| 11 | M162F,K167R,K170T,N174I,I175F | Cluster6 |
| 12 | A182T,A185N,V187N,G189A,N194T | Cluster7 |
| 13 | Y214L,L215M,T216K,I217A,D222Q | Cluster10,Cluster9 |
| 14 | S90L,M93T,N96K,V105N,N106A | Cluster1 |
| 15 | S116A,M119L,K120P,S125I,Y127Q | Cluster2 |
| 16 | K167R,S173F,N174I,I175F,F177V | Cluster6 |
| 17 | E128H,E130A,L132M,N134W,L136S,K138S | Cluster3 |
| 18 | S139D,S142I,A144L,I145S,S149M,L152V | Cluster3,Cluster4 |
| 19 | G202E,F203Y,H204P,P206S,E208L,E209D | Cluster8 |
| 20 | S224T,F226W,D227P,S235L,K236Q,Q238P,F239L | Cluster10 |
| 21 | I179T,D183G,D184I,R186E,I192F,R193K,R197H,F200Y | Cluster7 |
| 22 | E128H,E130A,L132M,G133S,N134W,L136S,N137K,K138S | Cluster3 |
| 23 | S139D,S141Q,S142I,A144L,I145S,H146Q,S149M,L152V | Cluster3,Cluster4 |
| 24 | S116A,M119L,K120P,S125I,Y127Q,K170T,N174I,F177V | Cluster2,Cluster6 |
| 25 | S116A,M119L,K120P,S125I,Y127Q,M162F,N174I,I175F | Cluster2,Cluster6 |
| 26 | S116A,M119L,K120P,S125I,Y127Q,K167R,N174I,I175F | Cluster2,Cluster6 |
| 27 | S116A,M119L,K120P,S125I,Y127Q,K167R,K170T,F177V | Cluster2,Cluster6 |
| 28 | S116A,M119L,K120P,S125I,Y127Q,K167R,K170T,I175F | Cluster2,Cluster6 |
| 29 | S116A,M119L,K120P,S125I,Y127Q,K170T,S173F,N174I,F177V | Cluster2,Cluster6 |
| 30 | S116A,M119L,K120P,S125I,Y127Q,M162F,K170T,N174I,F177V | Cluster2,Cluster6 |
| 31 | S116A,M119L,K120P,S125I,Y127Q,M162F,K170T,S173F,N174I | Cluster2,Cluster6 |
| 32 | S116A,M119L,K120P,S125I,Y127Q,M162F,K167R,K170T,N174I | Cluster2,Cluster6 |

| | | |
|---|---|---|
| **33** | S116A,M119L,K120P,S125I,Y127Q,K167R,K170T,N174I,I175F | Cluster2,Cluster6 |
| **34** | S116A,M119L,K120P,S125I,Y127Q,K167R,K170T,S173F,N174I,F177V | Cluster2,Cluster6 |
| **35** | S116A,M119L,K120P,S125I,Y127Q,K167R,K170T,S173F,I175F,F177V | Cluster2,Cluster6 |
| **36** | S116A,M119L,K120P,S125I,Y127Q,G133S,N137K,K167R,K170T,I175F | Cluster2,Cluster3,Cluster6 |
| **37** | N86H,E87D,F98H,Y99L,M101I,R102K,T107S,L109K,R111S,V112F,S113D | Cluster1 |
| **38** | S116A,M119L,K120P,S125I,Y127Q,K167R,K170T,S173F,F177V | Cluster2,Cluster6 |
| **39** | S116A,M119L,K120P,S125I,Y127Q,G133S,N137K,K167R,K170T,S173F,F177V | Cluster2,Cluster3,Cluster6 |
| **40** | S116A,M119L,K120P,S125I,Y127Q,E128H,G133S,N137K,K167R,K170T,F177V | Cluster2,Cluster3,Cluster6 |
| **41** | S116A,M119L,K120P,S125I,Y127Q,G133S,N137K,K167R,K170T,S173F,I175F,F177V | Cluster2,Cluster3,Cluster6 |
| **42** | N86H,E87D,S90L,F98H,Y99L,M101I,R102K,N106A,T107S,L109K,R111S,V112F,S113D | Cluster1 |
| **43** | S116A,M119L,K120P,S125I,Y127Q,E130A,L132M,N134W,L136S,K138S,K167R,K170T,F177V | Cluster2,Cluster3,Cluster6 |
| **44** | S116A,M119L,K120P,S125I,Y127Q,E128H,G133S,N137K,S141Q,M158L,K167R,K170T,F177V | Cluster2,Cluster3,Cluster4,Cluster5,Cluster6 |
| **45** | S116A,M119L,K120P,S125I,Y127Q,S139D,S141Q,S142I,A144L,I145S,H146Q,S149M,L152V,K167R,K170T,F177V | Cluster2,Cluster3,Cluster4,Cluster6 |
| **46** | S116A,M119L,K120P,S125I,Y127Q,E130A,L132M,N134W,L136S,K138S,S139D,S142I,A144L,I145S,H146Q,S149M,L152V,L157Y,K167R,K170T,F177V | Cluster2,Cluster3,Cluster4,Cluster5,Cluster6 |
| **47** | S116A,M119L,K120P,S125I,Y127Q,E128H,G133S,N137K,S139D,S141Q,S142I,A144L,I145S,H146Q,S149M,L152V,L157Y,M158L,K167R,K170T,F177V | Cluster2,Cluster3,Cluster4,Cluster5,Cluster6 |
| **48** | M93T,N96K,V105N,S116A,M119L,K120P,S125I,G133S,N137K,S141Q,H146Q,L157Y,M158L,M162F,K167R,K170T,N174I,I175F,A182T,A185N,V187N,G189A,N194T,V212L | Cluster1,Cluster2,Cluster3,Cluster4,Cluster5,Cluster6,Cluster7,Cluster8 |
| **49** | S116A,M119L,K120P,S125I,Y127Q,E128H,E130A,L132M,G133S,N134W,L136S,N137K,K138S,S139D,S141Q,S142I,A144L,I145S,H146Q,S149M,L152V,L157Y,M158L,K167R,K170T,F177V | Cluster2,Cluster3,Cluster4,Cluster5,Cluster6 |
| **50** | S90L,M93T,N96K,V105N,N106A,S116A,M119L,K120P,S125I,Y127Q,E128H,E130A,L132M,G133S,N134W,L136S,N137K,K138S,S139D,S141Q,S142I,A144L,I145S,H146Q,S149M,L152V,L157Y,M158L,K167R,K170T,F177V | Cluster1,Cluster2,Cluster3,Cluster4,Cluster5,Cluster6 |

| | | |
|---|---|---|
| **51** | S90L,M93T,N96K,V105N,N106A,S116A,M119L,K120P,S125I,Y127Q,E128H,E130A,L132M,G133S,N134W,L136S,N137K,K138S,S139D,S141Q,S142I,A144L,I145S,H146Q,S149M,L152V,L157Y,M158L,K167R,K170T,F177V,I179T | Cluster1,Cluster2,Cluster3,Cluster4,Cluster5,Cluster6,Cluster7 |
| **52** | S90L,M93T,N96K,V105N,N106A,S116A,M119L,K120P,S125I,Y127Q,E128H,E130A,L132M,G133S,N134W,L136S,N137K,K138S,S139D,S141Q,S142I,A144L,I145S,H146Q,S149M,L152V,L157Y,M158L,K167R,K170T,F177V,A182T | Cluster1,Cluster2,Cluster3,Cluster4,Cluster5,Cluster6,Cluster7 |
| **53** | S90L,M93T,N96K,V105N,N106A,S116A,M119L,K120P,S125I,Y127Q,E128H,E130A,L132M,G133S,N134W,L136S,N137K,K138S,S139D,S141Q,S142I,A144L,I145S,H146Q,S149M,L152V,L157Y,M158L,K167R,K170T,F177V,D183G | Cluster1,Cluster2,Cluster3,Cluster4,Cluster5,Cluster6,Cluster7 |
| **54** | S90L,M93T,N96K,V105N,N106A,S116A,M119L,K120P,S125I,Y127Q,E128H,E130A,L132M,G133S,N134W,L136S,N137K,K138S,S139D,S141Q,S142I,A144L,I145S,H146Q,S149M,L152V,L157Y,M158L,K167R,K170T,F177V,D184I | Cluster1,Cluster2,Cluster3,Cluster4,Cluster5,Cluster6,Cluster7 |
| **55** | S90L,M93T,N96K,V105N,N106A,S116A,M119L,K120P,S125I,Y127Q,E128H,E130A,L132M,G133S,N134W,L136S,N137K,K138S,S139D,S141Q,S142I,A144L,I145S,H146Q,S149M,L152V,L157Y,M158L,K167R,K170T,F177V,A185N | Cluster1,Cluster2,Cluster3,Cluster4,Cluster5,Cluster6,Cluster7 |
| **56** | S90L,M93T,N96K,V105N,N106A,S116A,M119L,K120P,S125I,Y127Q,E128H,E130A,L132M,G133S,N134W,L136S,N137K,K138S,S139D,S141Q,S142I,A144L,I145S,H146Q,S149M,L152V,L157Y,M158L,K167R,K170T,F177V,R186E | Cluster1,Cluster2,Cluster3,Cluster4,Cluster5,Cluster6,Cluster7 |
| **57** | S90L,M93T,N96K,V105N,N106A,S116A,M119L,K120P,S125I,Y127Q,E128H,E130A,L132M,G133S,N134W,L136S,N137K,K138S,S139D,S141Q,S142I,A144L,I145S,H146Q,S149M,L152V,L157Y,M158L,K167R,K170T,F177V,V187N | Cluster1,Cluster2,Cluster3,Cluster4,Cluster5,Cluster6,Cluster7 |
| **58** | S90L,M93T,N96K,V105N,N106A,S116A,M119L,K120P,S125I,Y127Q,E128H,E130A,L132M,G133S,N134W,L136S,N137K,K138S,S139D,S141Q,S142I,A144L,I145S,H146Q,S149M,L152V,L157Y,M158L,K167R,K170T,F177V,G189A | Cluster1,Cluster2,Cluster3,Cluster4,Cluster5,Cluster6,Cluster7 |
| **59** | S90L,M93T,N96K,V105N,N106A,S116A,M119L,K120P,S125I,Y127Q,E128H,E130A,L132M,G133S,N134W,L136S,N137K,K138S,S139D,S141Q,S142I,A144L,I145S,H146Q,S149M,L152V,L157Y,M158L,K167R,K170T,F177V,I192F | Cluster1,Cluster2,Cluster3,Cluster4,Cluster5,Cluster6,Cluster7 |
| **60** | S90L,M93T,N96K,V105N,N106A,S116A,M119L,K120P,S125I,Y127Q,E128H,E130A,L132M,G133S,N134W,L136S,N137K,K138S,S139D,S141Q,S142I,A144L,I145S,H146Q,S149M,L152V,L157Y,M158L,K167R,K170T,F177V,R193K | Cluster1,Cluster2,Cluster3,Cluster4,Cluster5,Cluster6,Cluster7 |
| **61** | S90L,M93T,N96K,V105N,N106A,S116A,M119L,K120P,S125I,Y127Q,E128H,E130A,L132M,G133S,N134W,L136S,N137K,K138S,S139D,S141Q,S142I,A144L,I145S,H146Q,S149M,L152V,L157Y,M158L,K167R,K170T,F177V,N194T | Cluster1,Cluster2,Cluster3,Cluster4,Cluster5,Cluster6,Cluster7 |
| **62** | S90L,M93T,N96K,V105N,N106A,S116A,M119L,K120P,S125I,Y127Q,E128H,E130A,L132M,G133S,N134W,L136S,N1 | Cluster1,Cluster2,Cluster3,Cluster4, |

| | | |
|---|---|---|
| | 37K,K138S,S139D,S141Q,S142I,A144L,I145S,H146Q,S149M,L152V,L157Y,M158L,K167R,K170T,F177V,R197H | Cluster5,Cluster6,Cluster7 |
| **63** | S90L,M93T,N96K,V105N,N106A,S116A,M119L,K120P,S125I,Y127Q,E128H,E130A,L132M,G133S,N134W,L136S,N137K,K138S,S139D,S141Q,S142I,A144L,I145S,H146Q,S149M,L152V,L157Y,M158L,K167R,K170T,F177V,F200Y | Cluster1,Cluster2,Cluster3,Cluster4,Cluster5,Cluster6,Cluster7 |
| **64** | S90L,M93T,N96K,V105N,N106A,S116A,M119L,K120P,S125I,Y127Q,E128H,E130A,L132M,G133S,N134W,L136S,N137K,K138S,S139D,S141Q,S142I,A144L,I145S,H146Q,S149M,L152V,L157Y,M158L,K167R,K170T,F177V,G202E | Cluster1,Cluster2,Cluster3,Cluster4,Cluster5,Cluster6,Cluster8 |
| **65** | S90L,M93T,N96K,V105N,N106A,S116A,M119L,K120P,S125I,Y127Q,E128H,E130A,L132M,G133S,N134W,L136S,N137K,K138S,S139D,S141Q,S142I,A144L,I145S,H146Q,S149M,L152V,L157Y,M158L,K167R,K170T,F177V,F203Y | Cluster1,Cluster2,Cluster3,Cluster4,Cluster5,Cluster6,Cluster8 |
| **66** | S90L,M93T,N96K,V105N,N106A,S116A,M119L,K120P,S125I,Y127Q,E128H,E130A,L132M,G133S,N134W,L136S,N137K,K138S,S139D,S141Q,S142I,A144L,I145S,H146Q,S149M,L152V,L157Y,M158L,K167R,K170T,F177V,H204P | Cluster1,Cluster2,Cluster3,Cluster4,Cluster5,Cluster6,Cluster8 |
| **67** | S90L,M93T,N96K,V105N,N106A,S116A,M119L,K120P,S125I,Y127Q,E128H,E130A,L132M,G133S,N134W,L136S,N137K,K138S,S139D,S141Q,S142I,A144L,I145S,H146Q,S149M,L152V,L157Y,M158L,K167R,K170T,F177V,P206S | Cluster1,Cluster2,Cluster3,Cluster4,Cluster5,Cluster6,Cluster8 |
| **68** | S90L,M93T,N96K,V105N,N106A,S116A,M119L,K120P,S125I,Y127Q,E128H,E130A,L132M,G133S,N134W,L136S,N137K,K138S,S139D,S141Q,S142I,A144L,I145S,H146Q,S149M,L152V,L157Y,M158L,K167R,K170T,F177V,E208L | Cluster1,Cluster2,Cluster3,Cluster4,Cluster5,Cluster6,Cluster8 |
| **69** | S90L,M93T,N96K,V105N,N106A,S116A,M119L,K120P,S125I,Y127Q,E128H,E130A,L132M,G133S,N134W,L136S,N137K,K138S,S139D,S141Q,S142I,A144L,I145S,H146Q,S149M,L152V,L157Y,M158L,K167R,K170T,F177V,E209D | Cluster1,Cluster2,Cluster3,Cluster4,Cluster5,Cluster6,Cluster8 |
| **70** | S90L,M93T,N96K,V105N,N106A,S116A,M119L,K120P,S125I,Y127Q,E128H,E130A,L132M,G133S,N134W,L136S,N137K,K138S,S139D,S141Q,S142I,A144L,I145S,H146Q,S149M,L152V,L157Y,M158L,K167R,K170T,F177V,V212L | Cluster1,Cluster2,Cluster3,Cluster4,Cluster5,Cluster6,Cluster8 |
| **71** | S90L,M93T,N96K,V105N,N106A,S116A,M119L,K120P,S125I,Y127Q,E128H,E130A,L132M,G133S,N134W,L136S,N137K,K138S,S139D,S141Q,S142I,A144L,I145S,H146Q,S149M,L152V,L157Y,M158L,K167R,K170T,F177V,Y214L | Cluster1,Cluster2,Cluster3,Cluster4,Cluster5,Cluster6,Cluster9 |
| **72** | S90L,M93T,N96K,V105N,N106A,S116A,M119L,K120P,S125I,Y127Q,E128H,E130A,L132M,G133S,N134W,L136S,N137K,K138S,S139D,S141Q,S142I,A144L,I145S,H146Q,S149M,L152V,L157Y,M158L,K167R,K170T,F177V,L215M | Cluster1,Cluster2,Cluster3,Cluster4,Cluster5,Cluster6,Cluster9 |
| **73** | S90L,M93T,N96K,V105N,N106A,S116A,M119L,K120P,S125I,Y127Q,E128H,E130A,L132M,G133S,N134W,L136S,N137K,K138S,S139D,S141Q,S142I,A144L,I145S,H146Q,S149M,L152V,L157Y,M158L,K167R,K170T,F177V,T216K | Cluster1,Cluster2,Cluster3,Cluster4,Cluster5,Cluster6,Cluster9 |

| | | |
|---|---|---|
| **74** | S90L,M93T,N96K,V105N,N106A,S116A,M119L,K120P,S125I,Y127Q,E128H,E130A,L132M,G133S,N134W,L136S,N137K,K138S,S139D,S141Q,S142I,A144L,I145S,H146Q,S149M,L152V,L157Y,M158L,K167R,K170T,F177V,I217A | Cluster1,Cluster2,Cluster3,Cluster4,Cluster5,Cluster6,Cluster9 |
| **75** | S90L,M93T,N96K,V105N,N106A,S116A,M119L,K120P,S125I,Y127Q,E128H,E130A,L132M,G133S,N134W,L136S,N137K,K138S,S139D,S141Q,S142I,A144L,I145S,H146Q,S149M,L152V,L157Y,M158L,K167R,K170T,F177V,D222Q | Cluster1,Cluster10,Cluster2,Cluster3,Cluster4,Cluster5,Cluster6 |
| **76** | S90L,M93T,N96K,V105N,N106A,S116A,M119L,K120P,S125I,Y127Q,E128H,E130A,L132M,G133S,N134W,L136S,N137K,K138S,S139D,S141Q,S142I,A144L,I145S,H146Q,S149M,L152V,L157Y,M158L,K167R,K170T,F177V,S224T | Cluster1,Cluster10,Cluster2,Cluster3,Cluster4,Cluster5,Cluster6 |
| **77** | S90L,M93T,N96K,V105N,N106A,S116A,M119L,K120P,S125I,Y127Q,E128H,E130A,L132M,G133S,N134W,L136S,N137K,K138S,S139D,S141Q,S142I,A144L,I145S,H146Q,S149M,L152V,L157Y,M158L,K167R,K170T,F177V,F226W | Cluster1,Cluster10,Cluster2,Cluster3,Cluster4,Cluster5,Cluster6 |
| **78** | S90L,M93T,N96K,V105N,N106A,S116A,M119L,K120P,S125I,Y127Q,E128H,E130A,L132M,G133S,N134W,L136S,N137K,K138S,S139D,S141Q,S142I,A144L,I145S,H146Q,S149M,L152V,L157Y,M158L,K167R,K170T,F177V,D227P | Cluster1,Cluster10,Cluster2,Cluster3,Cluster4,Cluster5,Cluster6 |
| **79** | S90L,M93T,N96K,V105N,N106A,S116A,M119L,K120P,S125I,Y127Q,E128H,E130A,L132M,G133S,N134W,L136S,N137K,K138S,S139D,S141Q,S142I,A144L,I145S,H146Q,S149M,L152V,L157Y,M158L,K167R,K170T,F177V,S235L | Cluster1,Cluster10,Cluster2,Cluster3,Cluster4,Cluster5,Cluster6 |
| **80** | S90L,M93T,N96K,V105N,N106A,S116A,M119L,K120P,S125I,Y127Q,E128H,E130A,L132M,G133S,N134W,L136S,N137K,K138S,S139D,S141Q,S142I,A144L,I145S,H146Q,S149M,L152V,L157Y,M158L,K167R,K170T,F177V,K236Q | Cluster1,Cluster10,Cluster2,Cluster3,Cluster4,Cluster5,Cluster6 |
| **81** | S90L,M93T,N96K,V105N,N106A,S116A,M119L,K120P,S125I,Y127Q,E128H,E130A,L132M,G133S,N134W,L136S,N137K,K138S,S139D,S141Q,S142I,A144L,I145S,H146Q,S149M,L152V,L157Y,M158L,K167R,K170T,F177V,Q238P | Cluster1,Cluster10,Cluster2,Cluster3,Cluster4,Cluster5,Cluster6 |
| **82** | S90L,M93T,N96K,V105N,N106A,S116A,M119L,K120P,S125I,Y127Q,E128H,E130A,L132M,G133S,N134W,L136S,N137K,K138S,S139D,S141Q,S142I,A144L,I145S,H146Q,S149M,L152V,L157Y,M158L,K167R,K170T,F177V,F239L | Cluster1,Cluster10,Cluster2,Cluster3,Cluster4,Cluster5,Cluster6 |
| **83** | N86H,E87D,F98H,Y99L,M101I,R102K,T107S,L109K,R111S,V112F,S113D,S116A,M119L,K120P,S125I,Y127Q,E128H,E130A,L132M,G133S,N134W,L136S,N137K,K138S,S139D,S141Q,S142I,A144L,I145S,H146Q,S149M,L152V,L157Y,M158L,K167R,K170T,F177V | Cluster1,Cluster2,Cluster3,Cluster4,Cluster5,Cluster6 |
| **84** | N86H,E87D,S90L,M93T,N96K,F98H,Y99L,M101I,R102K,V105N,N106A,T107S,L109K,R111S,V112F,S113D,S116A,M119L,K120P,S125I,Y127Q,E128H,E130A,L132M,G133S,N134W,L136S,N137K,K138S,S139D,S141Q,S142I,A144L,I145S,H146Q,S149M,L152V,L157Y,M158L,M162F,K167R,K170T,S173F,N174I,I175F,F177V,I179T,A182T,D183G,D18 | Cluster1,Cluster10,Cluster2,Cluster3,Cluster4,Cluster5,Cluster6,Cluster7,Cluster8,Cluster9 |

| | |
|---|---|
| | 4I,A185N,R186E,V187N,G189A,I192F,R193K,N194T,R197H,F200Y,G202E,F203Y,H204P,P206S,E208L,E209D,V212L,Y214L,L215M,T216K,I217A,D222Q,S224T,F226W,D227P,S235L,K236Q,Q238P,F239L |

**Table S3.** Statistics for cryoEM data collection, structure determination and model quality of wild-type V27 and V57 MACV GP1-hTfR1 complexes.

| Dataset | MACV GP1-hTfR1 | V27 GP1-hTfR1 | V57 GP1-hTfR1 |
|---|---|---|---|
| **Data collection and processing** | | | |
| **Microscope** | Titan Krios | Titan Krios | Titan Krios |
| **Voltage (keV)** | 300 | 300 | 300 |
| **Detector** | Falcon4i | Falcon4i | Falcon4i |
| **Nominal magnification** | 105,000 | 130,000 | 105,000 |
| **Data Acquisition Software** | EPU | EPU | EPU |
| **Electron dose ($e^{-}/Å^{2}$)** | 40 | 40 | 40 |
| **Pixel Size (Å)** | 1.2 | 0.95 | 1.2 |
| **Defocus range (µm)** | -0.5 to -1.5 | -0.5 to -1.5 | -0.5 to -1.5 |
| **Number of movies (#)** | 6,405 | 24,830 | 21,543 |
| **Number of particles** | 43,539 | 88,395 | 35,387 |
| **Symmetry imposed** | C2 | C2 | C1 |
| **Resolution (Å)** | 2.5 | 2.5 | 2.7 |
| **FSC threshold** | 0.143 | 0.143 | 0.143 |
| **Refinement** | | | |
| **Initial model used (PDB code)** | ***3KAS, 11ZB*** | ***3KAS, 11ZB*** | ***3KAS, 11ZB*** |
| **Non-hydrogen atoms** | 23466 | 23452 | 22181 |
| **Protein residues** | 2938 | 2936 | 2782 |
| **R.M.S. deviations** | | | |
| **Bond lengths (Å)** | 0.32 | 0.23 | 0.30 |
| **Bond angles (°)** | 0.35 | 0.35 | 0.42 |
| **Validation** | | | |
| **MolProbity score** | 1.50 | 1.59 | 1.62 |
| **Clashscore** | 4 | 4 | 6 |
| **Poor rotamers (%)** | 1 | 1 | 1 |
| **Ramachandran (%)** | | | |
| **Favored** | 96 | 95 | 96 |
| **Allowed** | 4 | 5 | 4 |
| **Disallowed** | 0.00 | 0 | 0 |
| **Fit to map ($CC_{mask}$)** | 0.85 | 0.84 | 0.80 |
| **Accession codes** | | | |
| **EMDB (maps)** | EMD-76440 | EMD-76443 | EMD-76444 |
| **PDB (model)** | 12HO | 12HS | 12HT |

### Figures

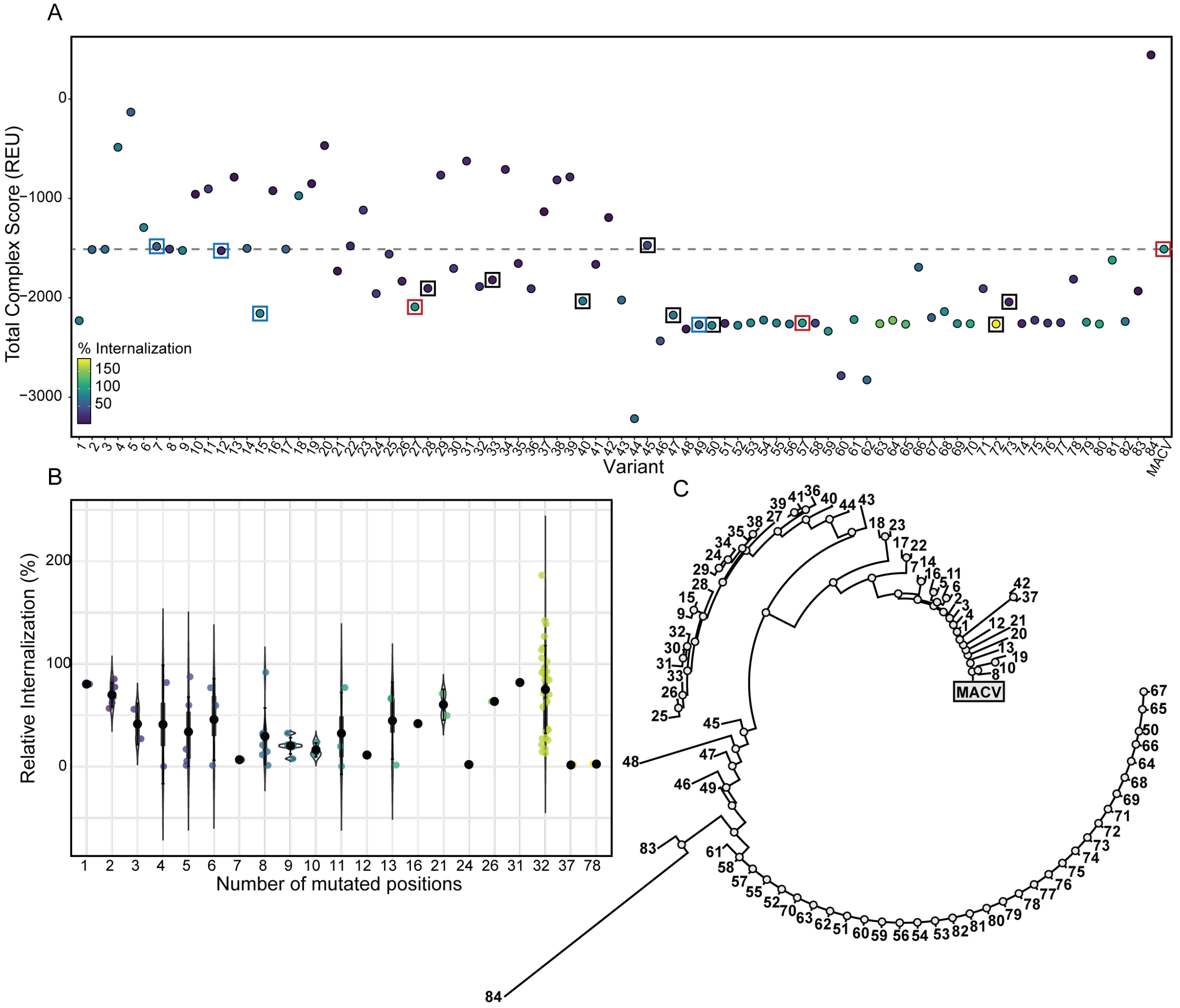

**Figure S1.** Selection and characterization of MACV/JUNV sequence variants. (A) Total complex scores predicted in Rosetta of each experimentally tested GP1 sequence variant as a full GPC model in complex with hTfR1$^{AD}$ after repacking. Red boxes denote sequences used for structural characterization, recombinant GP1 generation, and further biochemical characterization as refenced in Figures (2,3,4), blue boxes denote sequence variants used for further biochemical characterization and immunization studies as referenced in Figures (3,4,5), black boxes denote sequence variants sub-selected for further biochemical characterization as referenced in Figures (2,4). (B) Percent internalization of sequence variants relative to WT MACV GPC -bearing pseudoviruses plotted as a function of the number of mutations for each variant where the mean percent internalization is shown by a black point. (C) Phylogenetic tree constructed with ClustalW[27] after alignment of all experimentally tested GP1 sequence variants rooted to the MACV GP1 sequence.

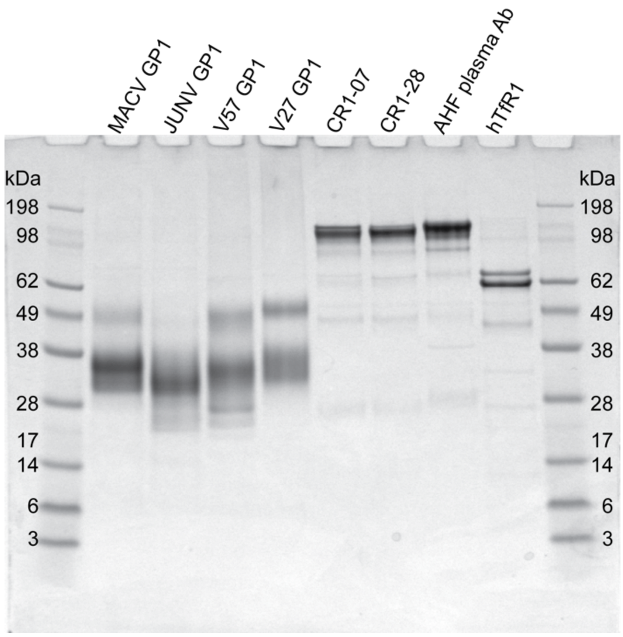

**Figure S2**. **Overall quality of recombinant NWM GP1 constructs, antibodies and hTfR1.** The overall purity and integrity of each recombinant protein was assessed by SDS-PAGE under non-reducing conditions. Lanes show glycosylated GP1 constructs of the appropriate molecular weight (20 kDa), properly assembled antibodies (150 kDa), and the hTfR1 ectodomain (~80 kDa) in complex with hTf (~75 kDa).

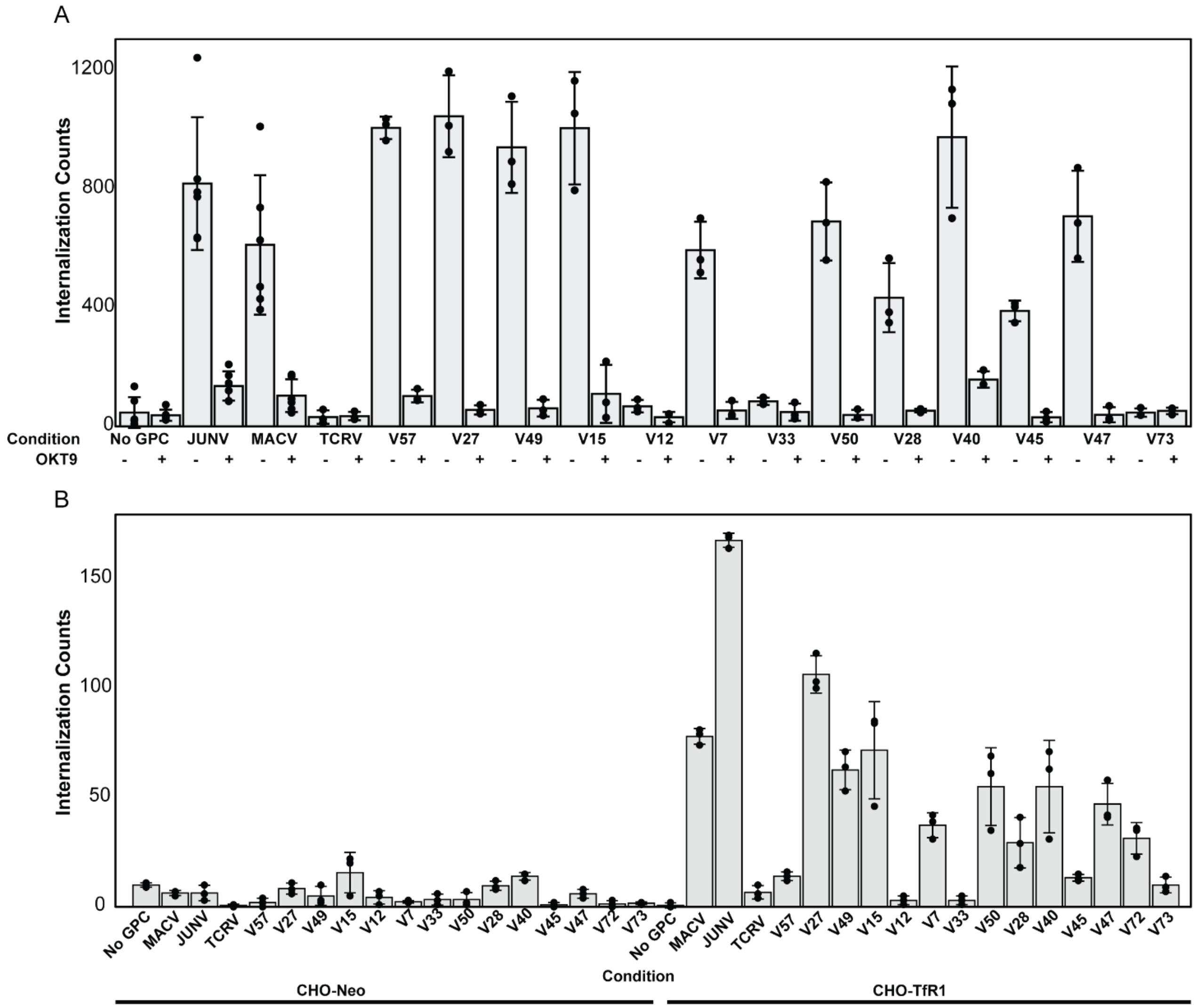

**Figure S3**. **TfR1-dependence in cellular internalization of a panel of select GPC variants.** (A) Internalization of select sequence variants into HEK293T cells treated with 100 μg/mL OKT9. (B) Internalization of select sequence variants into CHO-TfR1 and CHO-Neo cells.

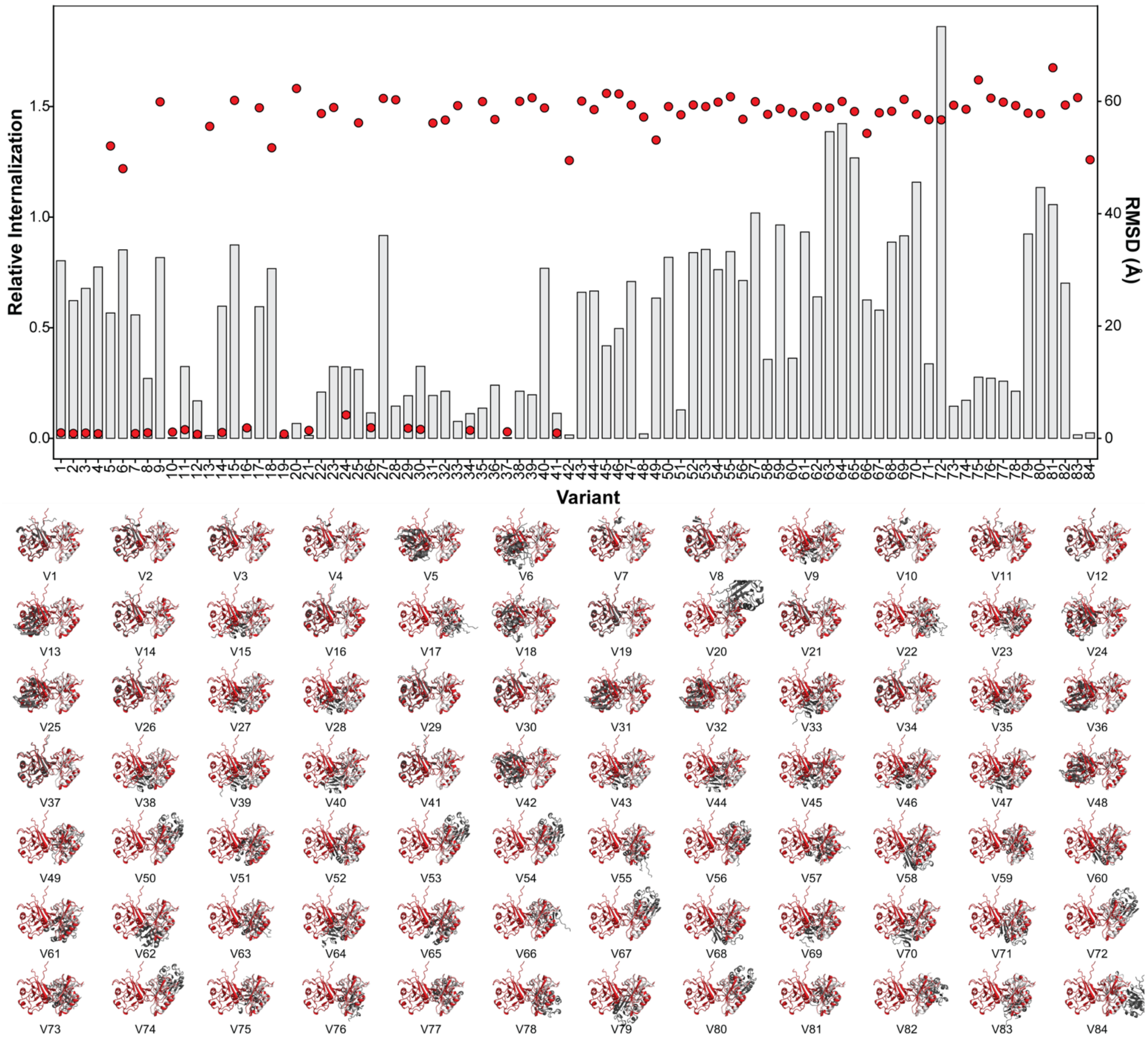

**Figure S4. AlphaFold prediction of hTfR1 apical domain binding by isolated GP1 constructs.** Average internalization of each GP1 sequence variant relative to WT MACV depicted by bar graph alongside the GP1 RMSD of AlphaFold predicted GP1 sequence variants in complex with $hTfR1^{AD}$ after alignment of each $hTfR1^{AD}$ domain to MACV-$hTfR1^{AD}$ and corresponding predicted structures of GP1 (white)- $hTfR1^{AD}$ (grey) overlaid with MACV GP1- $hTfR1^{AD}$ (red).

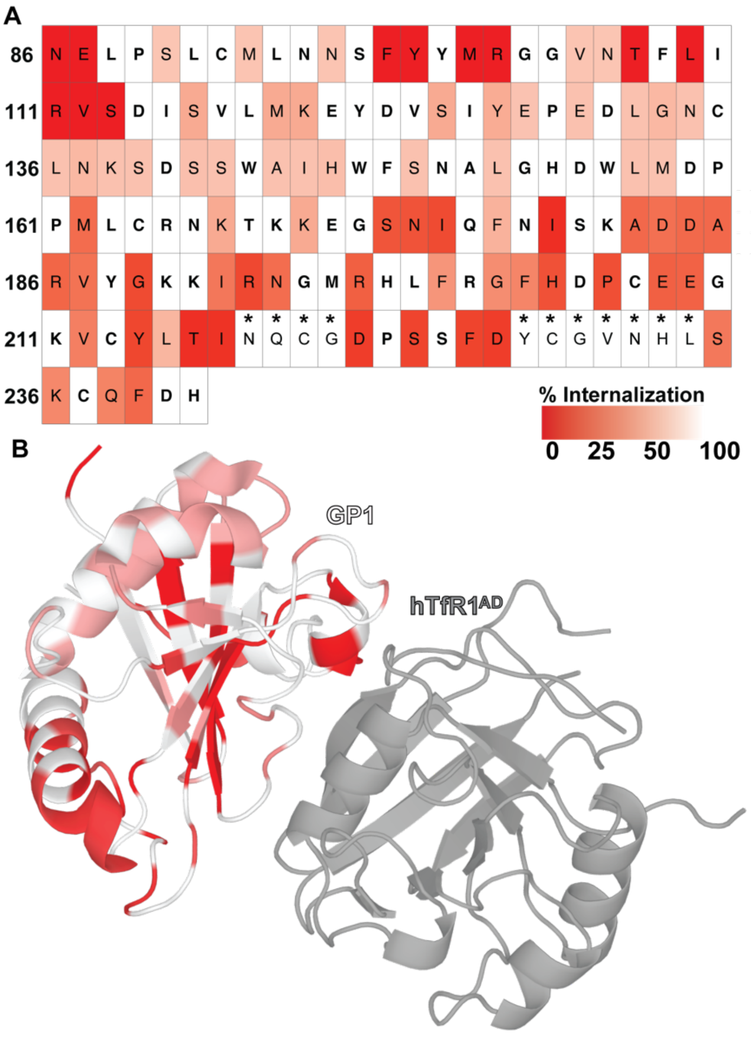

**Figure S5. Average percent internalization across sequence variants at each site relative to WT MACV.** (A) Percent internalization shown by sequence where non-mutated/conserved residue positions between MACV and JUNV are bolded and asterisks denote MACV insertion sites. (B) Percent internalization colored on the GP1-hTfR1$^{AD}$ derived from PDB:3KAS[20].

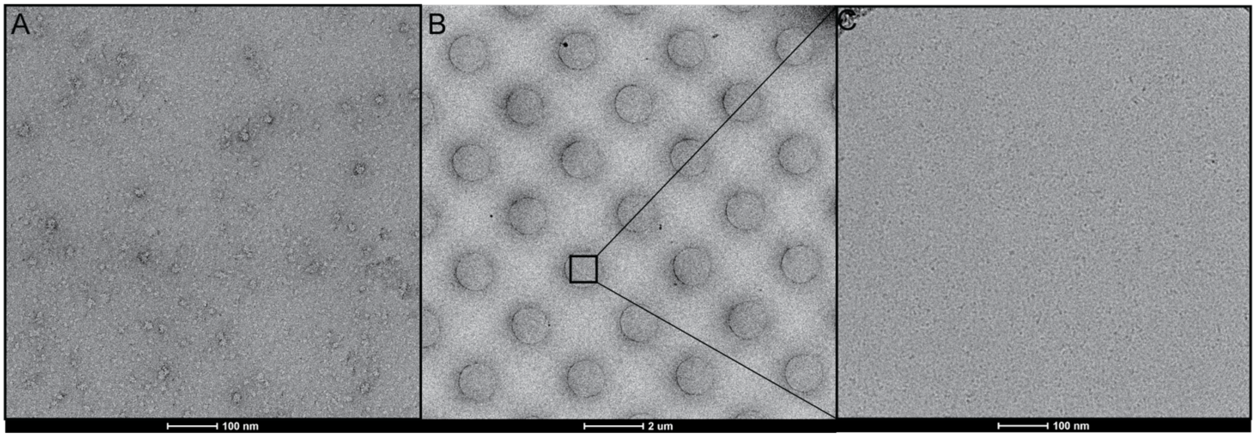

**Figures S6**. **Preparation of GP1-hTfR1 complexes for cryoEM analysis.** (A) Representative negative stain EM micrograph of MACV GP1-hTfR1; scale bar: 100 nm. (B) Representative cryoEM micrograph of MACV GP1-hTfR1 at low magnification; scale bar: 2 μm. B) Representative cryoEM micrograph of MACV GP1-hTfR1 at high magnification; scale bar: 100nm.

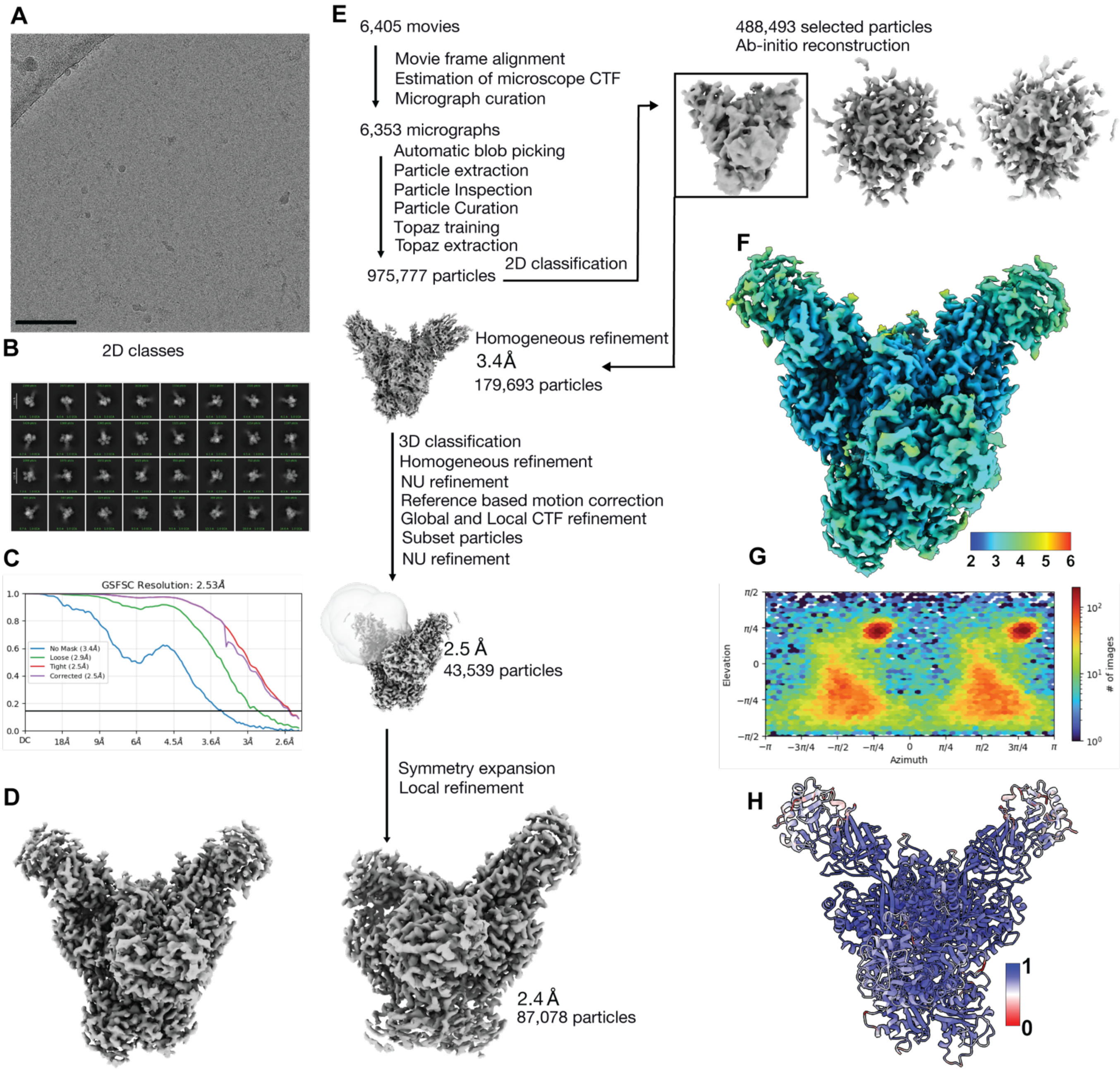

**Figures S7**. **cryoEM data processing of the MACV GP1-hTfR1 complex.** (A) Representative cryoEM micrograph of MACV GP1-hTfR1; scale bar: 100 nm. (B) 2D classes of the selected MACV GP1-hTfR1 particles. (C) Gold-standard Fourier shell correlation curve with the 0.143 threshold indicated by a horizontal black line. (D) Sharpened final map at 2.5 Å. (E) cryoEM data processing workflow. CTF = contrast transfer function. (F) Local resolution map for the MACV GP1-hTfR1 complex. (G) The particle orientation density plot calculated using cryoSPARC is shown below the local resolution map. (H) MACV GP1-hTfR1 model colored by Q-score values, which indicate the correlation between the built atomic model and the corresponding cryoEM map[28].

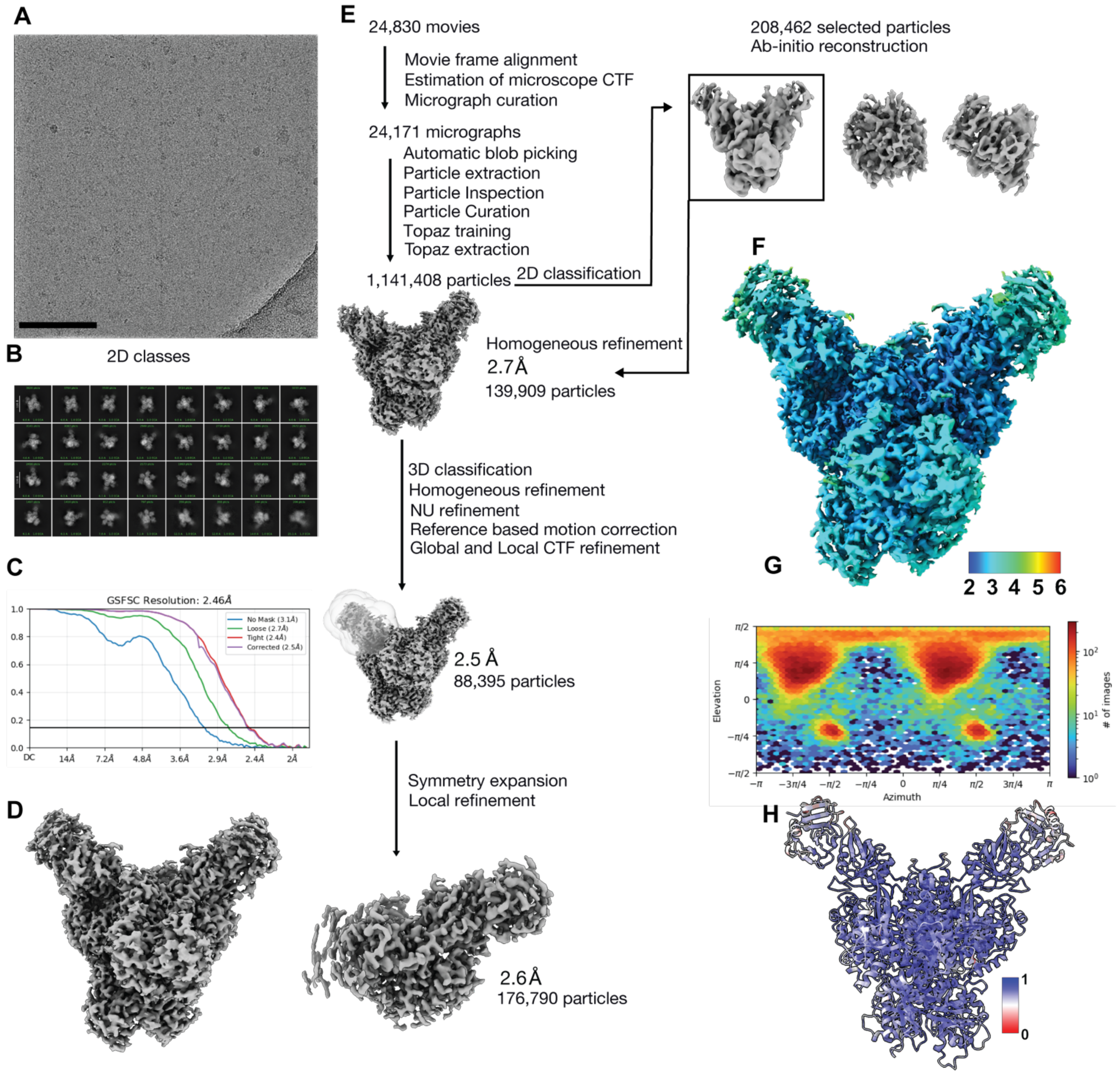

**Figures S8**. **cryoEM data processing of the V27 GP1-hTfR1 complex.** (A) Representative cryoEM micrograph of V27 GP1-hTfR1; scale bar: 100 nm. (B) 2D classes of the selected V27 GP1-hTfR1 particles. (C) Gold-standard Fourier shell correlation curve with the 0.143 threshold indicated by a horizontal black line. (D) Sharpened final map at 2.5 Å. (E) cryoEM data processing workflow. CTF = contrast transfer function. (F) Local resolution map for the V27 GP1-hTfR1 complex. (G) The particle orientation density plot calculated using cryoSPARC is shown below the local resolution map. (H) V27 GP1-hTfR1 model colored by Q-score values, which indicate the correlation between the built atomic model and the corresponding cryoEM map[28].

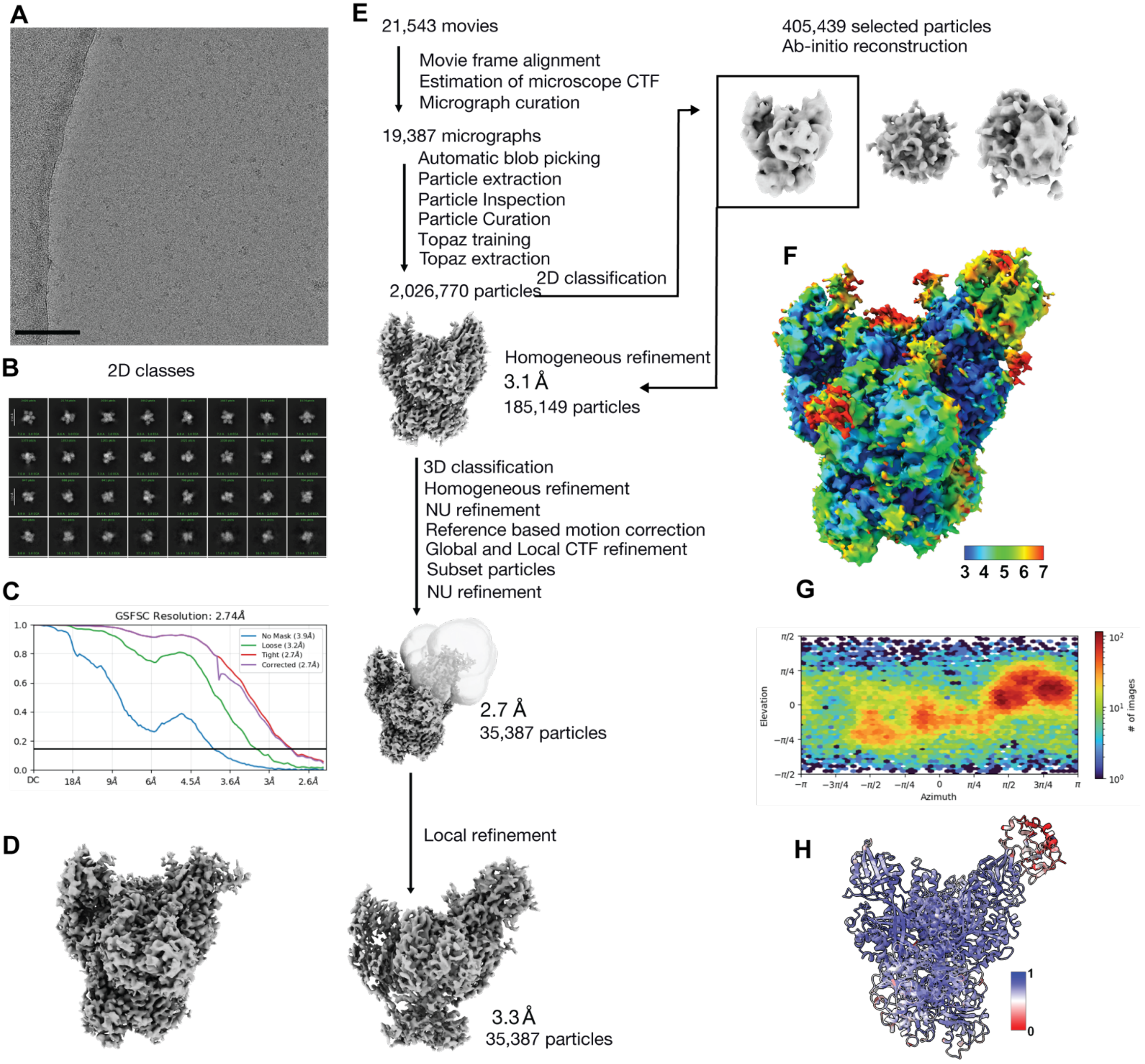

**Figures S9**. **cryoEM data processing of the V57 GP1-hTfR1 complex.** (A) Representative cryoEM micrograph of V57 GP1-hTfR1; scale bar: 100 nm. (B) 2D classes of the selected V57 GP1-hTfR1 particles. (C) Gold-standard Fourier shell correlation curve with the 0.143 threshold indicated by a horizontal black line. (D) Sharpened final map at 3.2 Å. (E) cryoEM data processing workflow. CTF = contrast transfer function. (F) Local resolution map for the V57 GP1-hTfR1 complex. (G) The particle orientation density plot calculated using cryoSPARC is shown below the local resolution map. (H) V57 GP1-hTfR1 model colored by Q-score values, which indicate the correlation between the docked atomic model and the corresponding cryoEM map[28].
